# Sensitive detection of DNA contamination in tumor samples via microhaplotypes

**DOI:** 10.1101/2020.12.18.423488

**Authors:** Brett Whitty, John F. Thompson

## Abstract

Low levels of sample contamination with other human DNAs can have disastrous effects on the accurate identification of somatic variation in tumor samples. Detection of sample contamination in DNA is often based on low frequency variants that indicate if more than a single source of DNA is present. This strategy works with standard DNA samples but can be problematic in solid tumor FFPE samples because there are often huge variations in allele frequency (AF) due to copy number changes arising from gains and losses across the genome. The variable AFs make detection of contamination challenging. To avoid this, we counted microhaplotypes to assess sample contamination. Microhaplotypes are sets of variants on the same sequencing read that can be unambiguously phased. Instead of measuring AF, the number of microhaplotypes is determined. Contamination detection becomes based on fundamental genomic properties, linkage disequilibrium (LD) and the diploid nature of human DNA, rather than variant frequencies. We optimized microhaplotype panel content and selected 164 SNV sets located in regions already being sequenced within a cancer panel. Thus, contamination detection uses existing sequence data. LD data from the 1000 Genomes Project is used to make the panel ancestry agnostic, providing the same sensitivity for contamination detection with samples from individuals of African, East Asian, and European ancestry. Detection of 1% contamination with no matching normal sample is possible. The methods described here can also be extended to other DNA mixtures such as forensic and non-invasive prenatal testing samples where DNA mixes can be similarly detected. The microhaplotype method allows sensitive detection of DNA contamination in FFPE tumor and other samples when deep coverage with Illumina or other high accuracy NGS is used.

## Introduction

In many applications, DNA contamination is not a significant issue because small amounts of extraneous DNA do not affect the experimental outcome. When examining DNA from a single species, it is easy to filter out sequences from other species but within-species contamination is more challenging. When analyzing germ-line variants, a substantial fraction of human DNA from other sources is required to affect variant calls because non-reference calls will be approximately 0%, 50% or 100%. For example, concordance between variant calls made by SNP chips versus pure NGS for high depth samples with 5% contamination fell only from 99.7% to 99.2% (1). However, even with low-level DNA contamination, there can be a significant impact on some applications like detecting low frequency somatic variants in tumor DNA samples or deconvoluting complex mixtures in forensic samples. Clinical diagnostic samples are presumed to be pure when they are tested, but there is always a concern that unexpected contamination could affect results and escape standard detection methods. Knowledge of whether a sample is contaminated with another and, if so, the extent of that contamination, is critical for assessing whether somatic variant calls can be trusted.

Contamination detection is often carried out by measuring variable lengths of short tandem repeats (STRs) or by genotyping/sequencing single-nucleotide variants (SNVs). When unusual frequencies are observed, the degree of contamination is estimated by quantitating the frequency of the extra variants (2-5). However, accurate estimation becomes difficult at low variant frequency because the true signal can be obscured by the competing noise arising from technical issues like STR stutter or genotyping/sequencing errors. This is especially problematic with analog measurement systems that are often used with STR, genotyping, and classical sequencing methods. In contrast, next-generation sequencing (NGS) methods produce a digital signal, allowing greater sensitivity and accuracy at low contamination levels.

In addition to the method described here, there are informatic methods that make use of different types of NGS read data to model whether contamination is likely. These methods are often tied to particular data and/or sample types so may not extrapolate well to other systems like clinical test samples. For example, ContEst (2) is highly sensitive but uses both whole genome and microarray data or matched normal samples. This has recently been replaced by CalculateContamination that relaxes copy number and matched sample requirements. Verifybamid (6) can be used with just NGS data, but its accuracy is dependent on large numbers of SNVs and knowledge of population SNV frequencies and cannot be used with DNA with large copy number changes. It is also computationally intensive; but, VerifyBamID2 was recently developed to address these issues (7). Conpair (8) is much less computationally intensive but still requires large numbers of SNVs as well as tumor-normal pairs. It can be used with exomes or large panels but not smaller panels. Sensitive detection of circulating transplant DNA has been achieved with genome-wide SNVs (9) but this method is hampered when there is copy number variation as is often found in tumor samples. Sehn et al. (10) have taken advantage of closely spaced SNVs that are not in perfect LD, known as microhaplotypes (MH), to assess DNA contamination to a level of 5%. Microhaplotypes can be defined as multiple variants on the same sequence read such that the phasing of the variants is known and the haplotype can be unambiguously determined.

Testing clinical tumor samples for somatic variation is complicated by the presence of large regions of the tumor genome that may be highly amplified or deleted. When diagnostic samples have large fractions of the genome with altered copy number, the allelic frequencies (AFs) of SNVs can vary substantially and change from the typical 50-50 AF to much more extreme values. These altered allele ratios can overlap the range of AFs for what is typically seen with contamination and thus a sample that is 100% pure can mistakenly appear to be significantly contaminated if only AF is examined. To avoid these problems, we have optimized an MH counting method that is independent of genotype AF for determining contamination status with tumor DNA and can be carried out with no matching normal sample.

For individuals with a normal set of chromosomes, each autosomal region of any size has two sets of alleles. Those alleles may be the same (homozygous) or different (heterozygous); but, under normal circumstances with a few specific exceptions, there should never be more than two haplotypes in a genomic region in any individual. When DNA from two individuals is mixed, accidentally or intentionally, there is a possibility for 1-4 MHs in each region. MHs have been used for forensic purposes to detect contamination and identify individuals (11-17), to detect transplant DNA in a host background and contamination in tumor samples (18), and to determine ancestry (19-23). Use of MHs for routine detection and estimation of contamination in clinical FFPE tumor samples is a powerful application of these methods.

## Materials and Methods

### Sequencing Samples

849 de-identified lung tumor samples and DNA from lung tumors were obtained from several commercial and academic providers. These included BioIVT, Conversant Bio, Cureline, Duke, Folio Bio, Fundacio Institute, Indivumed, iSpecimen, NCI, and Proteogenex. Samples were received either as FFPE material or extracted DNA. If necessary, extraction of DNA from FFPE specimens was carried out using standard kits. DNA was prepared for sequencing using hybridization capture with panels directed primarily at exons in >500 genes implicated in cancer (see https://www.personalgenome.com/assets/resources/elio-tissue-complete-brochure.pdf for a listing of genes). For some genes, intronic and promoter DNA was also targeted. DNA was sequenced using an Illumina NextSeq500. The PGDx elio™ tissue complete assay was used for sample preparation and analysis (24, 25). 15 samples were sequenced on standard runs. For a run to pass, at total of >90 GB sequence with >75% of reads at Q30 or above was required. For an individual sample to pass, >90% of targeted regions were required to have >100x median coverage. Individual reads were analyzed as described (26) with a base quality filter applied to only include bases with a reported Phred quality score >30. Because erroneous reads could artificially inflate the number of 3^rd^/4^th^ MHs, MH calculations included 3^rd^ MHs only if they surpassed a minimum fraction of total calls. We usually set this threshold at >0.2% based on the typical error rate for Illumina NGS (∼0.1%). Higher values can be used to reduce noise but that is accompanied by a loss in sensitivity.

Each accepted read covering the positions of all SNVs for each MH set was binned into the appropriate haplotype and counted using Samtools mpileup. After read alignment, mpileup was applied to the first region, the second region, and, if present, the third region. mpileup reads are then combined and separated by basecalls. Each resultant set of base calls (MHs) is then counted to determine MH frequency.

### *In silico* read mixing

Artificially contaminated samples were made by *in silico* mixing of reads at defined contamination levels. Contamination levels of x% were generated by combining randomly selected 100-x% reads from the sample with x% random reads from the contaminant and running the resultant reads through the standard analysis pipeline. It was found that the same results were achieved if only the reads mapping to the MH regions being examined were included in the mixing. These analyses involved much less data so ran on the pipeline much faster. Thus, later experiments used only the truncated data set to speed analysis and allow more conditions to be tested.

## Results

### DNA samples and MAF-based contamination detection

849 FFPE tumor DNA samples were examined, but most studies focused on the 51 samples listed in Table 1. All samples that passed QC were amenable to contamination testing, but the number of samples analyzed for mixing had to be limited due to the high compute requirements for analyzing the many *in silico* mixtures. The mixing samples were chosen based on being the most apparently pure and equally representing three of the five geographic ancestral groups described by the 1000 genomes project (African, East Asian, European). DNA quality was highly variable which primarily impacted mixing studies when two samples of much different quality were mixed.

**Table 1.** Microhaplotypes and Coverage for Pure Samples. Performance of the 164 SNV set panel is shown with 51 pure samples with known ancestries. Comparisons include the median and average coverage, the number of SNV sets with 1-4 MHs, and the median frequency of 3^rd^ MHs among individuals of African (Afri), East Asian (EaAs), and European (Euro) ancestry as defined by the 1000 Genomes project. The number and median 3^rd^ MH frequency for MHs with frequency >0.2% is shown. ND=Not Done; NA=Not Applicable

| Sample | M/<br>F | Ancestry | Number of MHs |  |  |  |  |  |  |  |  |  |
| --- | --- | --- | --- | --- | --- | --- | --- | --- | --- | --- | --- | --- |
|  |  |  | 1 | 2 | 3 | 4 | Med<br>3rd<br>MH | # 3<sup>rd</sup><br>MHs<br>>0.002 | Med<br>3<sup>rd</sup><br>>0.002 | # | Med<br>Cov | Ave<br>Cov |
| AATF748T |  | Afri | 52 | 87 | 21 | 3 | 0.0033 | 18 | 0.0081 | 163 | 313 | 620 |
| AATF094T | F | Afri | 36 | 90 | 36 | 0 | 0.0027 | 29 | 0.0031 | 162 | 421 | 533 |
| AATF115T | F | Afri | 67 | 82 | 12 | 0 | 0.0027 | 7 | 0.0059 | 161 | 202 | 286 |
| AATF066T | F | Afri | 48 | 92 | 21 | 0 | 0.0041 | 17 | 0.0044 | 161 | 305 | 337 |
| AATF117T | F | Afri | 37 | ## | 22 | 1 | 0.0053 | 18 | 0.0065 | 163 | 439 | 605 |
| AATF498T | F | Afri | 68 | 76 | 19 | 0 | 0.0018 | 8 | 0.0037 | 163 | 674 | 959 |
| AATF308T | M | Afri | 42 | 96 | 24 | 0 | 0.0022 | 14 | 0.0033 | 162 | 531 | 683 |
| AATF562T |  | Afri | 46 | ## | 14 | 1 | 0.0021 | 8 | 0.0029 | 163 | 574 | 706 |
| AATF486T | F | Afri | 39 | 88 | 36 | 0 | 0.0023 | 21 | 0.0034 | 163 | 709 | 924 |
| PGRD0454T | M | Afri | 58 | 90 | 14 | 0 | 0.0018 | 6 | 0.0027 | 162 | 797 | 930 |
| AATF218T | F | Afri | 42 | 88 | 33 | 1 | 0.0029 | 24 | 0.0034 | 164 | 405 | 555 |
| AATF778T |  | Afri | 60 | 91 | 11 | 1 | 0.0023 | 7 | 0.0055 | 163 | 462 | 520 |
| AATF219T | M | Afri | 33 | 95 | 34 | 1 | 0.0022 | 19 | 0.0039 | 163 | 553 | 668 |
| AATF217T | M | Afri | 57 | 90 | 15 | 0 | 0.0018 | 5 | 0.0029 | 162 | 609 | 996 |
| AATF073T | F | EaAs | 74 | 78 | 12 | 0 | 0.0021 | 7 | 0.0028 | 164 | 415 | 726 |
| AATF503T | F | EaAs | 62 | 76 | 24 | 1 | 0.0031 | 16 | 0.0037 | 163 | 426 | 545 |
| AATF504T | M | EaAs | 70 | 84 | 8 | 1 | 0.0042 | 9 | 0.0042 | 163 | 517 | 758 |
| AATF027T | M | EaAs | 58 | 92 | 13 | 1 | 0.0024 | 8 | 0.0052 | 164 | 331 | 443 |
| AATF012T | F | EaAs | 66 | 91 | 7 | 0 | 0.0029 | 5 | 0.0050 | 164 | 542 | 826 |
| AATF029T | M | EaAs | 42 | 74 | 46 | 1 | 0.0033 | 37 | 0.0035 | 163 | 379 | 590 |
| AATF024T | F | EaAs | 31 | 82 | 48 | 2 | 0.0037 | 39 | 0.0051 | 163 | 452 | 724 |
| AATF023T | M | EaAs | 33 | 91 | 36 | 3 | 0.0025 | 26 | 0.0028 | 163 | 500 | 765 |
| AATF022T | M | EaAs | 42 | 92 | 30 | 0 | 0.0023 | 19 | 0.0046 | 164 | 510 | 768 |
| AATF021T | M | EaAs | 56 | 78 | 29 | 0 | 0.0027 | 19 | 0.0041 | 163 | 528 | 792 |
| AATF733T | F | EaAs | 53 | 93 | 17 | 0 | 0.0047 | 12 | 0.0086 | 163 | 170 | 257 |
| AATF595T | F | EaAs | 53 | 84 | 27 | 0 | 0.0038 | 19 | 0.0098 | 164 | 201 | 390 |
| AATF742T | F | EaAs | 51 | ## | 10 | 0 | 0.0020 | 5 | 0.0128 | 163 | 210 | 352 |
| AATF734T | M | EaAs | 60 | 89 | 13 | 1 | 0.0029 | 11 | 0.0032 | 163 | 344 | 431 |
| AATF594T |  | EaAs | 64 | 92 | 7 | 0 | 0.0028 | 5 | 0.0030 | 163 | 390 | 533 |
| AATF735T | M | EaAs | 68 | 88 | 6 | 1 | 0.0025 | 5 | 0.0027 | 163 | 399 | 614 |
| AATF597T | F | EaAs | 54 | 94 | 14 | 1 | 0.0021 | 8 | 0.0062 | 163 | 399 | 502 |
| AATF606T | F | EaAs | 62 | 85 | 15 | 0 | 0.0023 | 10 | 0.0029 | 162 | 443 | 659 |
| AATF601T | F | EaAs | 58 | 85 | 20 | 0 | 0.0022 | 11 | 0.0030 | 163 | 457 | 570 |
| AATF599T | F | EaAs | 58 | 84 | 21 | 1 | 0.0025 | 15 | 0.0042 | 164 | 487 | 648 |
| AATF600T | F | EaAs | 52 | 94 | 16 | 1 | 0.0024 | 10 | 0.0028 | 163 | 605 | 752 |
| AATF417T | M | Euro | 67 | 90 | 5 | 1 | 0.0142 | 5 | 0.0176 | 163 | 187 | 239 |
| AATF391T | M | Euro | 47 | 95 | 20 | 1 | 0.0019 | 10 | 0.0023 | 163 | 476 | 754 |
| AATF375T | M | Euro | 34 | 97 | 32 | 1 | 0.0052 | 30 | 0.0056 | 164 | 305 | 463 |
| AATF088T | F | Euro | 41 | 99 | 21 | 1 | 0.0029 | 16 | 0.0038 | 162 | 288 | 406 |
| AATF389T | F | Euro | 39 | 94 | 30 | 1 | 0.0021 | 16 | 0.0030 | 164 | 446 | 741 |
| AATF567T | M | Euro | 68 | 84 | 10 | 0 | 0.0035 | 8 | 0.0053 | 162 | 312 | 704 |
| AATF107T | M | Euro | 51 | 98 | 13 | 1 | 0.0034 | 9 | 0.0066 | 163 | 445 | 518 |
| AATF079T | F | Euro | 58 | 94 | 10 | 0 | 0.0016 | 4 | 0.0041 | 162 | 494 | 641 |
| AATF496T | M | Euro | 62 | 94 | 8 | 0 | 0.0018 | 2 | 0.0322 | 164 | 601 | 674 |
| AATF036T | M | Euro | 44 | 97 | 20 | 2 | 0.0029 | 15 | 0.0030 | 163 | 538 | 603 |
| AATF042T | M | Euro | 39 | 93 | 31 | 1 | 0.0020 | 16 | 0.0027 | 164 | 613 | 747 |
| AATF039T | M | Euro | 49 | ## | 11 | 0 | 0.0020 | 5 | 0.0035 | 162 | 693 | 842 |
| AATF038T | F | Euro | 50 | 94 | 16 | 2 | 0.0015 | 7 | 0.0031 | 162 | 673 | 789 |
| AATF731T | F | Euro | 59 | 87 | 17 | 1 | 0.0024 | 12 | 0.0027 | 164 | 352 | 495 |
| AATF378T | F | Mix | 52 | 94 | 17 | 1 | 0.0028 | 12 | 0.0042 | 164 | 418 | 707 |
| AATF097T | F | Mix | 35 | 98 | 30 | 0 | 0.0035 | 22 | 0.0042 | 163 | 416 | 486 |
| Median |  |  | 52 | 91 | 17 | 1 | 0.0025 | 11 | 0.0038 | 163 | 445 | 641 |
| Mean |  |  | 52 | 90 | 20 | 1 | 0.0030 | 13 | 0.0052 | 163 | 450 | 623 |

Prior to using MHs, we had assessed DNA contamination based on finding low level germline SNVs that were assumed to come from contaminating DNA. However, FFPE tumor samples can have high levels of copy number variation that can have a serious impact on germline SNV AFs as shown in Fig 1. For 10 FFPE tumor samples, all variants in the VCF with a dbSNP number were examined. Having an assigned dbSNP number was used as a surrogate for variation that is most likely real since it has been observed previously. Variants were sorted by MAF into 10% bins. Most samples had the expected distribution where >80% of the variants had 0-10% MAF with this low-level variation arising from sequencing errors, FFPE artifacts or somatic variants. In addition, there are also many with 40-50% MAF arising from heterozygous germline variants. However, three samples had much more widely distributed MAFs including one sample (AATF748T) where >20% of the variants had MAFs between 20 and 30%. These are germline variants as confirmed by matching normal DNA samples that were also sequenced for these samples. When we used a method for detecting DNA contamination based on low frequency germline variants, these samples were discarded based on supposed high contamination levels when, in fact, they are close to 100% pure based on MH analysis. The “contamination” signal is actually copy number variation.

**Figure 1.**
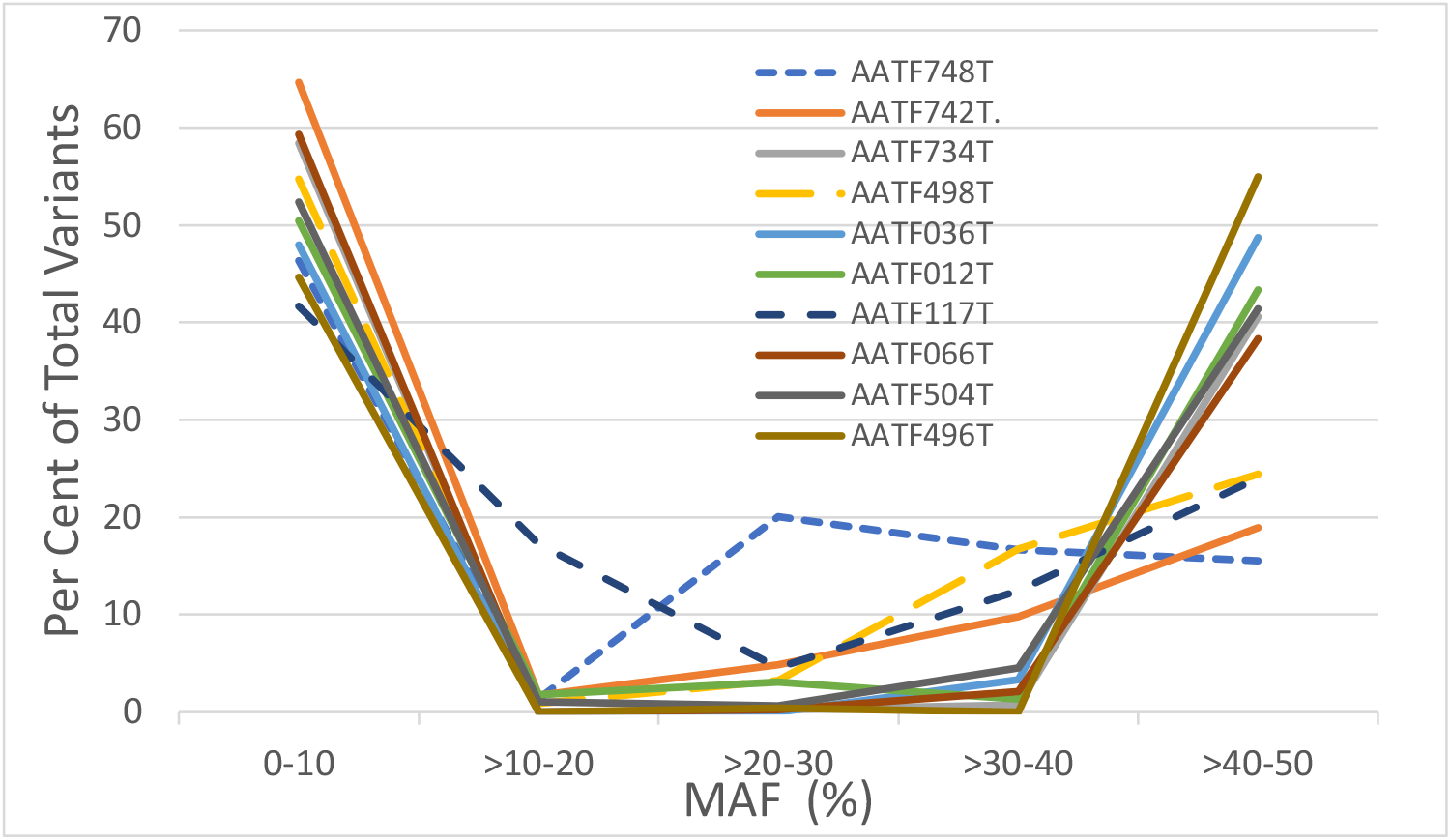
MAF for 10 FFPE tumor samples. All variants with dbSNP identifiers were binned by AF for the less common allele. Bins were 10% ranges. These samples had 348-561 such variants in the VCF with the vast majority being germline as confirmed by matching normal samples. The percentage of variants in each bin is shown. Additional information about these samples is in Table 1.

### Choice of variant sets for microhaplotype analysis

There are many MHs that have been characterized in the literature, and their usefulness for distinguishing individuals and ancestries is well documented (13, 20, 23). However, most of the well-characterized MHs are not in regions chosen for commercial cancer panel sequencing. In addition, the criteria for selecting MHs for discriminating individuals and ancestry are not the same as for detecting contamination where the goal is not distinguishing individuals but maximizing the number of MHs with potentially >2 versions. Since high depth sequencing is required for sensitive contamination detection using this approach, regions already targeted for analysis were examined so that additional, non-productive sequencing in MH regions outside of the pre-existing panel could be avoided. For gene panel sequencing, this means that literature MH data is of limited value. Each novel MH set we chose was tested for behavior across samples and ancestries. For these experiments, we have used a >500 gene, 2.23 Mb cancer panel directed against FFPE tumor DNA (https://www.personalgenome.com/assets/resources/elio-tissue-complete-brochure.pdf), but any pre-existing panel can be examined in a similar manner to identify useful SNV sets. This cancer panel included regions outside of exons and these regions were also inspected for potentially informative SNV sets.

Candidate MH sets were chosen manually, but these steps can be automated if many or larger panels are desired. The criteria for inclusion in the initial set of variants to be tested included maximizing the frequency of 3^rd^ MHs while minimizing the likelihood of sequencing errors in those variants. This was achieved by avoiding regions with higher inherent sequencing error rates such as insertions, deletions, and variants within homopolymeric regions and other repetitive elements. The regions targeted for cancer panel sequencing were examined in the gnomAD database (gnomad.broadinstitute.org/; 27) for multiple SNVs within 200 bp of the targeted regions and of each other and with 3-97% AF. The desired final AF (5-95%) was higher, but we wanted to ensure that MHs from populations underrepresented in gnomAD could still be identified. The gnomAD database, at the time initially queried, had a higher proportion of individuals of European ancestry (∼60%) than individuals with other geographic ancestries. Because of this, variation in European ancestry individuals had a larger impact on the initial choice of SNVs than variation in other ancestries. Thus, there is a possibility that samples from different ancestral backgrounds could respond differently when not filtered based on ancestry. Once the candidate MH sets were identified, they were balanced for ancestry so that detection of contamination in all groups would behave similarly.

Variants not in a segmental duplication or a low confidence region as defined by gnomAD were examined for LD in the 1000 Genomes Project samples (ldlink.nci.nih.gov/?tab=ldhap;28). This site provides data on the frequency of each haplotype in each ancestry. Pairs and triplets of SNVs with at least three haplotypes and the 3^rd^ and greater haplotypes having a total frequency of >5% in any individual ancestral grouping in the 1000 genomes project were advanced for evaluation in real samples sequenced as described. SNV sets with median coverage >250x for all relevant variants on individual reads were examined further. SNV sets with high frequency 3^rd^/4^th^ MHs in many purportedly pure samples were eliminated because they might produce high noise relative to signal. With these samples, SNV sets with more than 5 samples with >2 MHs were eliminated (26 of 264 evaluated SNV sets). SNV sets in close physical proximity to each other in high LD would provide duplicate information. LD was considered too high if the haplotype frequency was unchanged when the SNVs were combined and examined together. SNV sets with shorter homopolymers, higher coverage, and fewer 3^rd^/4^th^ MHs in pure samples were included in the panel when high LD alternatives were considered. SNV sets that passed these initial evaluations are shown in Supplementary Table 1. In addition to the genomic coordinates and dbSNP identifiers, the frequency of MHs is shown for the five major ancestry groupings defined by 1000 Genomes as well as the median coverage for the relevant SNVs across 849 samples, the number of times the SNV set generated more than 2 MHs in “pure” samples, and the DNA length separating the SNVs.

Once these criteria were met, SNV sets were balanced for ancestral frequency. While it would have been desirable to use all population groupings from 1000 Genomes in this analysis, there were not enough SNV sets to allow balancing of all five major ancestry groupings so only data from African, East Asian, and European ancestry were assessed to achieve balanced variation. SNV sets that adversely affected ancestry balance were eliminated. The balanced panel of 164 SNV sets is shown in Supplementary Table 1. The average frequency of 3^rd^/4^th^ MHs was 13.9% for each of the three balanced ancestries.

### Assessing contamination and estimating levels

In order to determine how best to detect contamination, *in silico* mixing of reads from individual pure samples was carried out so that the precise level of contamination would be known and the sensitivity and accuracy of the MH assay could be determined. The initial samples used for establishing contamination methods need to be as close to 100% pure as possible for maximizing sensitivity. 51 samples were chosen for *in silico* mixing experiments to examine this in more detail. Samples with self-declared ancestry were chosen to achieve ancestral diversity and the fewest number of >2 MH sets. The performance of 51 reportedly pure samples from individuals of known, self-declared ancestry is shown in Table 1.

The self-declared ancestry of samples was confirmed using Principle Component Analysis (PCA) of the SNV sets based on 1000 Genome MH frequencies. The self-declared and PCA-determined ancestries of all samples agreed. However, with some samples, the self-declared ancestry was Asian and the PCA-determined ancestry was South Asian rather than East Asian. For analysis of individuals who listed Asian ancestry, only samples with PCA-confirmed East Asian ancestry were used in analyses. In addition to diverse ancestral histories, samples were selected for mixing experiments based on having the fewest SNV sets with >2 MHs.

For these samples, the mean number of 3^rd^/4^th^ MHs is over 20. Because these samples are supposed to be pure, the 3^rd^/4^th^ MHs are likely often due to artifactual false calls that arise due to the high coverage depth for many SNV sets or FFPE-induced errors. This is supported by the mean 3^rd^ MH frequency which is only 0.003 for the panel. To minimize the number of artifactual 3^rd^ MHs, we set a lower limit of 0.2% for counting 3^rd^/4^th^ MHs based on the typical Illumina error rate of ∼0.1%. Using a threshold of >0.2% eliminates over 1/3 of 3^rd^ MHs in these pure samples, most of which are likely to be artifactual. When this threshold is employed, the median 3^rd^ MH frequency increases but remains less than 0.6% for all samples with >20 3^rd^ MHs. The need for a minimum threshold for the number of 3^rd^/4^th^ MHs is highlighted by the three examples from these 51 samples where the median 3^rd^ MH frequency is >0.01. All three have five or fewer 3^rd^ MHs, suggesting that outlier artifactual MHs are driving the median frequency when few real 3^rd^ MHs are present. Based on this concern, we instituted a minimum threshold of 25 SNV sets with >2 MHs with frequency >0.2%. 25 SNV sets with >2 MHs is not intended to distinguish contaminated from pure samples but rather ensure that the 3^rd^ MH frequency is more reliable. 5 of the 51 pure DNAs in Table 1 have > 25 3^rd^/4^th^ MHs (>0.2% minimum threshold for inclusion as a 3^rd^ MH). This threshold should never eliminate a truly contaminated DNA but is intended only to ensure there are enough SNV sets to establish a reliable 3^rd^ MH median frequency. The low rate of 3^rd^/4^th^ MHs across pure samples support the concept that, at least at the sites analyzed, NGS and FFPE errors occur at a very low rate. All *in silico* mixtures tested at the 0.5% contamination level using the samples in Table 1 had more than 25 MHs. As seen in Fig 2, the average value for the number of 3^rd^ MHs among all ancestries with 1% contamination is >45 with this panel (range 29-69). Thus, this threshold should not eliminate any samples contaminated at >0.5% and allows the 3^rd^ MH frequency value to be more reliable.

**Figure 2.**
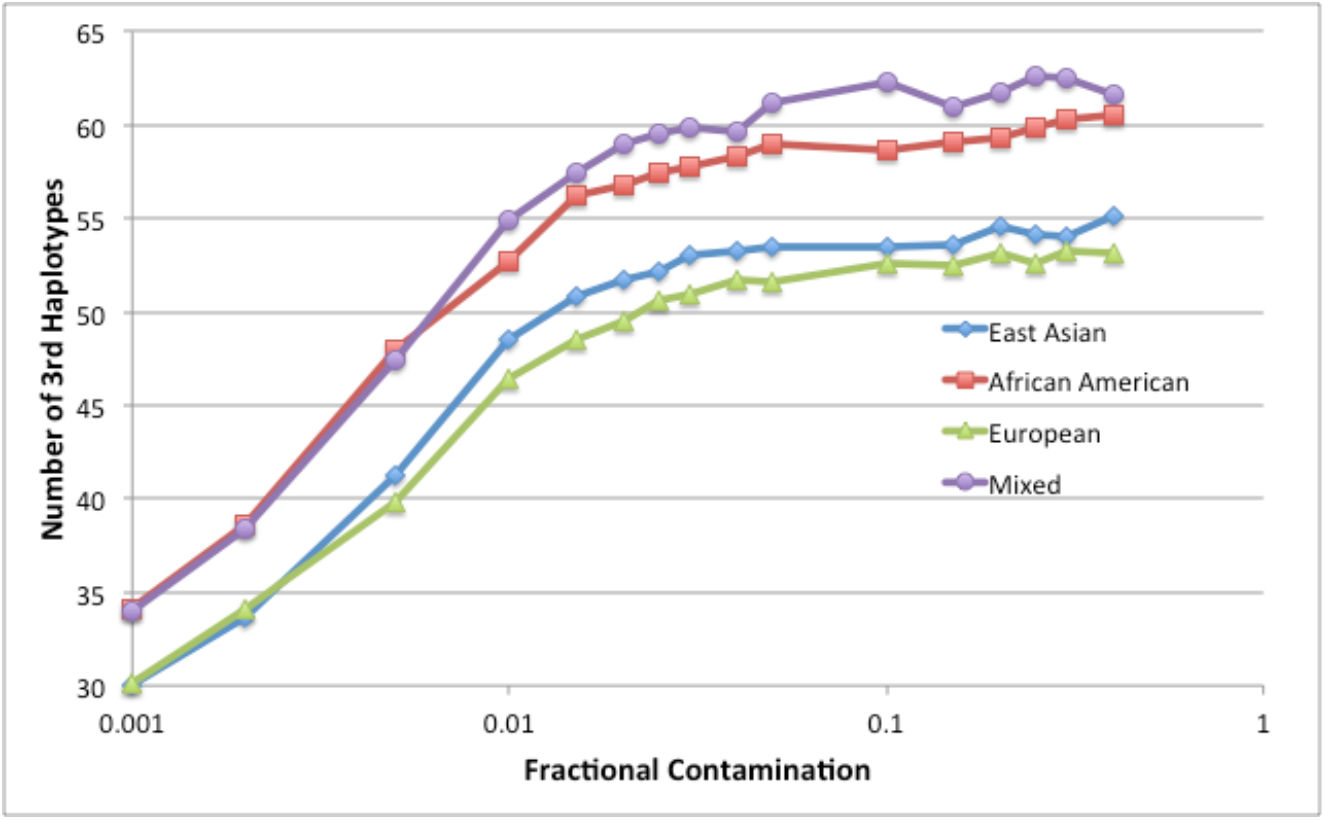
Number of SNV sets with >2 MHs for individual samples. The number of 3^rd^ MHs for pairs of individuals of East Asian, African American, and European ancestry is shown for mixes of 0.1 – 40%. Mixing results from pairs of individuals with different ancestry is included as “Mixed”, all on a log scale.

Precise levels of read mixing can be generated *in silico* but the functional level of contamination was found to often differ significantly from the known mixing proportions. This issue was particularly obvious when samples with divergent DNA qualities were mixed. The longer, higher quality DNA appeared to have a higher concentration than what had been added to the mix.This arises because longer reads map better and are more likely to contain all the variants being examined and thus this DNA appears to be relatively more frequent than the shorter, more degraded DNA. This is consistent with previous results that showed DNA with reduced extraction yields tend to be lower quality and more susceptible to contamination (10). To minimize the impact of this effect, all DNA mixes were examined in a pairwise fashion where one sample was the primary sample and the second was the contaminant. These roles were then switched and the results averaged. This allowed an over-contamination with one sample set to be partially mitigated by the paired under-contamination of the opposite sample set.

15 DNA sets from each ancestry were mixed as sample/contaminant pairs with averaged results shown in Fig 2. As expected, the East Asian fraction of >2 MHs was nearly identical to the European results. The African American samples that we examined had a higher frequency of >2 MHs than the other groupings, but nearly identical to the samples that were mixed across ancestries. This is predicted by 1000 Genomes data. Their subgroupings of African ancestry included seven different sub-populations, only one of which was US-based. When the African American subgrouping is compared to the other African populations, it has a higher 3^rd^/4^th^ MH frequency (0.155 vs 0.139, Supplementary Table 1). The higher frequency in our population reflects the fact our African ancestry samples are primarily from African American individuals.

Even at the lowest tested contamination level, 0.1%, the mean number of 3^rd^/4^th^ MHs is over 30 compared to 13 (range 2-39) for the pure samples without mixing (Table 1). At 1% contamination, the mean number of 3^rd^/4^th^ MHs is over 45 for all groupings with a range for individual mixes of 29-69. Because a small number of 3^rd^/4^th^ MHs can cause a median values to be impacted greatly by outliers, we have set a minimum threshold for the number of 3^rd^/4^th^ MHs before a contamination level is calculated. Based on these data, 25 is set as the minimum number before a sample is considered potentially contaminated because it is less than all samples mixed at 1% and high enough to minimize the impact of individual outliers.

While the number of 3^rd^ MHs is relevant for assessing whether a sample is potentially contaminated, it is less useful for estimating the level of contamination as it reaches a maximum around 2% contamination with these coverage levels. In contrast, median 3^rd^ MH frequency changes as a function of contamination level so is more useful in this regard.

The *in silico* mixing studies provide an empirical calibration, but it is also useful to understand the theoretical basis for those findings. As shown in Fig 3, the expected frequency of the contaminating 3^rd^ MH depends on the nature of the starting and incoming genotypes. Some combinations will generate no 3^rd^ MH, others will generate a 3^rd^ MH with a frequency half the level of incoming contaminant, and others will generate a genotype that is the same frequency as the contaminant. Since most variant combinations will be at the 50% level, examination of the median value for the 3^rd^ MH should yield a value that is half of the true contamination level.

**Figure 3.**
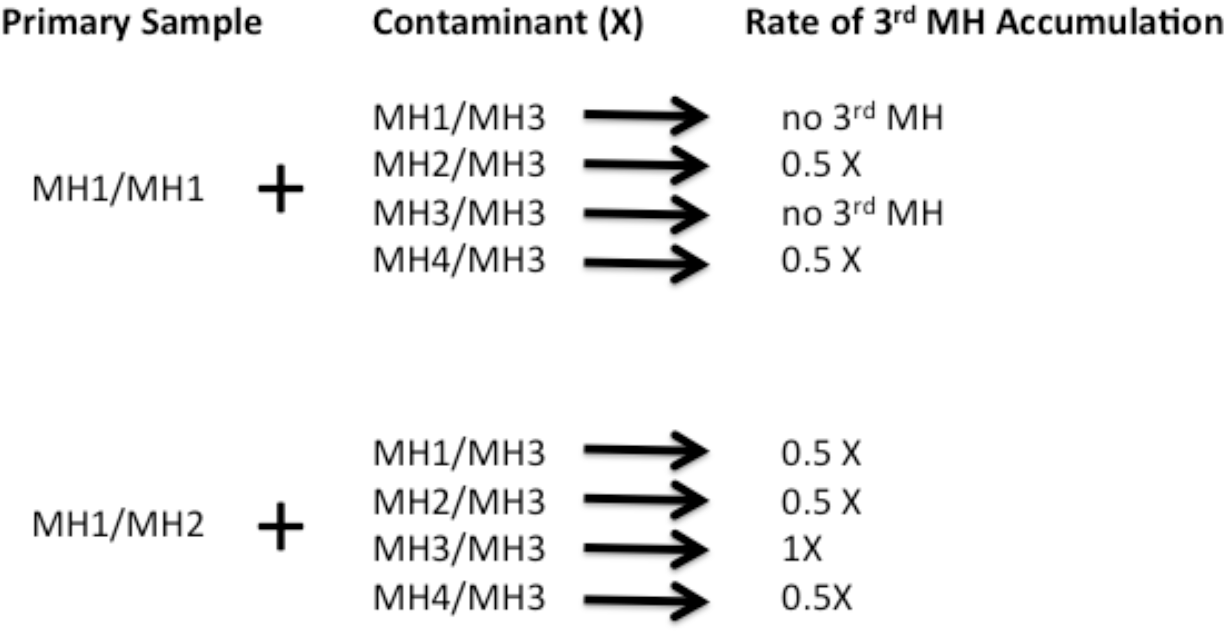
Expected frequencies for 3^rd^ MHs based on genotype. A primary sample can be either homozygous for one MH or heterozygous for two versions. For an incoming contaminating sample with at least one third MH, there are four different possibilities for each starting primary sample as shown. Most of these generate a contamination signal at half the frequency of the starting DNA. When the four alleles are all different, the 3^rd^ MH comes in at half the frequency while the combined 3^rd^/4^th^ frequency is 1x. If there is a 4^th^ MH, it should have the same frequency as the 3^rd^ MH as long as there are no technical artifacts.

The primary limitations on the sensitivity of contamination detection are the practical sequencing depths for the SNV sets in question and the inherent error rates for sequencing. The higher the sensitivity desired, the greater sequencing depth required. The usefulness of added depth is limited by sequencing error rates that are typically on the order of 0.1% for Illumina. It is possible to employ various error correction methods to improve sensitivity, but the need for that is dependent on the sensitivity required for the application. In addition, there is no value in sequencing a sample more deeply than the starting number of input molecules, which can be a limitation in some situations. Based on these considerations, we have aimed for detection of contamination at least as low as 1% in these samples.

As shown in Fig 4, the median 3^rd^ MH level is independent of ancestry with this panel. All three ancestries and mixed samples behave nearly identically. These data are combined in Supplementary Table 2. 15 sample/contaminant pairs were tested for each ancestry with the same number of each mixed ancestry pairs. When all samples are aggregated, 40% of samples with 0.5% contamination were detected. If the threshold is set at 1%, all contaminated samples were detected. The detection threshold can be set wherever required based on the detection needs and the likely impact of contamination on final results. However, the core limitations of sequencing accuracy, coverage depth, and input DNA molecules combine to ultimately limit sensitivity.

**Figure 4.**
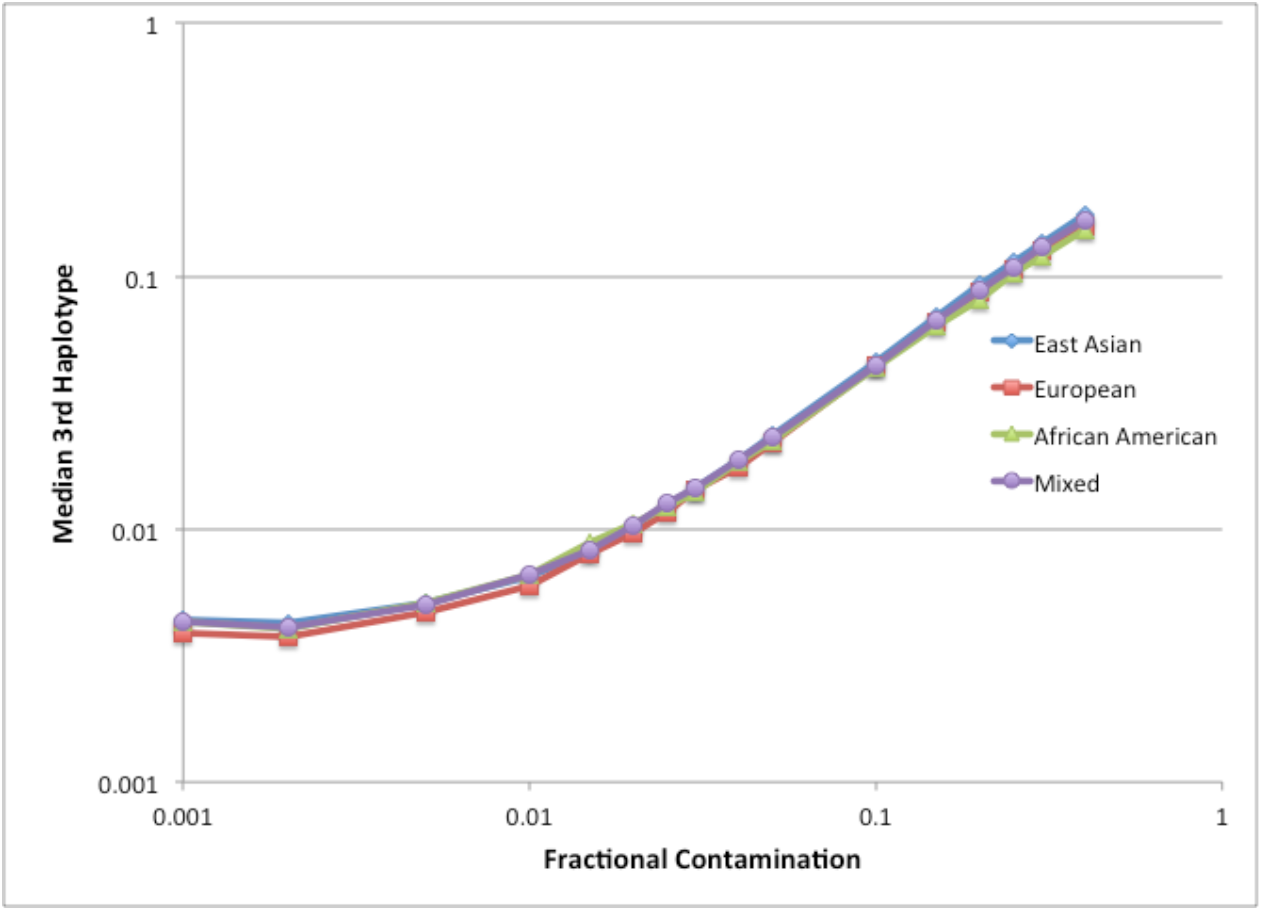
Contamination Detection as a function of ancestry. The median frequency of 3^rd^ MHs for pairs of individuals of East Asian, African American, and European ancestry is shown. Results from pairs of individuals with different ancestry is included as “Mixed”, all on a log scale for mixing levels of 0.1 – 40%.

In addition to detecting contamination, it is also possible to use this method to identify sources of contamination. A partial set of genotypes for the contaminating DNA can be generated from the MH data and then matched with samples that have been tested previously or contemporaneously to determine if they are the source of contamination. In a contaminated sample, MHs with 1 or 4 different haplotypes can be assigned their correct genotypes directly. MHs with 2 or 3 different haplotypes can generally be assigned a partial or complete genotype based on the frequency of the individual MHs. The frequency of the 2^nd^/3^rd^ haplotypes will provide at least one and sometimes two genotypes corresponding to the contaminant. Comparison of the known and partial genotypes with other samples allows them to be ruled in or out as potential sources of contamination. This procedure has been applied to multiple contaminated samples to successfully determine the source in real clinical laboratory situations. Often, the contamination source can be shown not to arise from any samples known to have been handled in the laboratory so the contamination must have originated prior to sample receipt.

## Discussion

Cross-contamination of DNA samples is a well-known issue among clinical testing laboratories and hence standard clinical testing guidelines generally suggest that precautions should be taken to be prevent it. Two recent guidelines for somatic variant testing (29, 30) recommend checking for handling-induced contamination and, suggest some possible methods for detecting contamination (6, 10, 31). However, none of the methods mentioned is sensitive enough to ensure that somatic variants (32) can be distinguished from sample contamination. Thus, simple, sensitive methods for reliable detection of contamination are needed.

The degree of DNA contamination that causes problems varies depending on the application. In some cases, the impact of contamination on results is obvious. With other applications, signals arising from contamination can be easily confused with real signals. With diagnosis of somatic variants in cancer, this is especially a problem because variant identification is attempted at and below the very limits of the NGS technology. Being able to separate the tumor signal from any contamination noise is critical for the proper diagnosis of tumors. Because tumor samples often have highly unusual copy number patterns, use of simple genotype frequencies can cause errors in contaminant detection. In our experience, many FFPE samples that had been determined to be contaminated using raw genotype frequencies were found to be pure by MH analysis (Fig 1). In contrast to FFPE tumor samples, circulating-free DNA (cfDNA) purified from the plasma of cancer patients is not as prone to the same issues. Most cfDNA purified from plasma is from normal cells so typical Mendelian ratios are observed with variants. MH analysis in such samples is still valuable for sensitive contamination detection, but such samples are not as likely to generate false positives as DNA from FFPE tumor samples where copy number variation is much more significant because of the high tumor DNA content.

For most purposes, the approximation that the percent contamination is twice the median frequency for the 3^rd^ MH is sufficient. This relationship arises from the likelihood that a 3^rd^ MH is most likely heterozygous so is present on only one of the two incoming alleles. The uncertainty around this probability introduces the potential for multiple sources of error and those should be considered if a more exact measurement is necessary. At low coverage, two confounding factors arise. First, low coverage leads to high AF variability due to the stochastic nature of read accumulation in poorly covered SNV sets. Only SNV sets with 3^rd^ MHs are included in the median calculation rather than all SNV sets so, at low coverage, AF inflation occurs among the variants that are counted. The observed AF is higher than reality because there are not enough reads to generate the proper AF value. By requiring a large number of 3^rd^ MHs prior to calling a sample contaminated, this effect is mitigated. Superimposed on this effect is the counting of homozygous 3^rd^ MHs before heterozygous 3^rd^ MHs as coverage increases. If coverage is deep, this is not an issue but, at low coverage, homozygous variants will appear first because they are being double counted. Further complicating the relationship is the presence of low frequency technical artifacts and the copy number issues that can occur with cancer samples. The copy number variations will make SNV sets appear at different fractional rates depending on whether the 3^rd^ MH is under- or over-represented. The more SNV sets used in the calculations, the more likely it is that these effects can be averaged out.

The MH approach is perfectly suited for use when somatic variants are sought because such studies use very high coverage in order to detect rare variants. The sensitivity of contamination detection is limited by the coverage of the regions used in the analysis. Other panel-based and exome NGS approaches can be made more reliable by using similar contamination detection methods. A typical whole genome approach with less than 50x coverage would not be amenable to sensitive detection using individual MH sets because the low coverage would prevent the observation of infrequent 3^rd^ MHs. To overcome this, combinations of MH sets could be used. The simplest approach would be to combine SNV sets that are in high LD. Other methods for pooling SNV sets could also be used.

The theoretical limitation on the sensitivity of the MH assay is the number of DNA molecules available for study. The assay can only be as sensitive as the amount of input DNA allows. In addition to that limitation (which is significant for some samples), there are also technology limitations. When looking at a pair of SNVs, either one could be called incorrectly resulting in false 3^rd^ MHs. If panels with 3 or more SNVs in close proximity were available, this would minimize such errors. However, there are other potential errors introduced in NGS processing. It is not unusual for barcodes used for multiplexing samples in the same run to be contaminated at 1% or more, resulting in misassignment of reads to samples. Index hopping and chimeric PCR reads can also affect sensitivity (33, 34).

The same techniques used for these cancer samples can also be used with forensic samples to detect contributors to a sample. Unlike tumor samples, copy number is not an issue. Like tumor samples, the major and minor contributors to a sample may have different DNA qualities that can affect the accuracy of MH frequencies, but this is less important in forensic situations. Panels designed with a larger number of >4 potential MHs would be appropriate when forensic samples are being examined to clarify the number of contributors.

Another useful application of the MH methods is for detection of CNV anomalies in fetal DNA in Non-invasive prenatal testing (NIPT). MH methods are already in use for paternity testing (35-38). For NIPT, panels can be designed to focus on genomic regions with the most likely copy number variation (e. g. chr 5, 13, 18, or 21) and the frequency of 3^rd^ MHs in targeted regions compared to the rest of the genome. In genomic regions where the mother is heterozygous and the father’s contribution is a different allele, there will be a low frequency 3^rd^ MH.

The methods described here highlight the factors that are important for examining DNA mixtures in a variety of contexts. The use of MHs in these applications requires different properties than other uses (39). Most early studies with MHs focus on distinguishing individuals and assigning ancestry. The MH properties required in those situations are different with a need for MHs specific for particular populations. With MHs used for contamination detection, it is preferable to have MHs that are common to all populations so that detection is uniform across individuals. The methods described herein allow the appropriate choice of MHs leading to the ability to detect low levels of secondary DNA, which can be valuable in many applications.

## Abbreviations

AdAm: Admixed American
AF: Allele Frequency
AfAm: African American
Afri: African
cfDNA: circulating free DNA
EaAs: East Asian
Euro: European
FFPE: Formalin-fixed Paraffin-embedded
FP: False Positive
LD: Linkage Disequilibrium
MH: Microhaplotype
NGS: Next-generation Sequencing
NIPT: Non-invasive Prenatal Testing
PCA: Principle Component Analysis
SNV: Single Nucleotide Variant
SoAs: South Asian
STR: Short Tandem Repeat

---

**Supplementary Table 1.** Detailed characterization of MH sets. The genomic locations of all SNV sets tested are shown in column 1. Median coverage in 849 samples and the number of instances of more than 2 MHs are shown in the next columns. The distance between SNVs and the dbSNP identifiers are listed in the next columns. The final columns show the sum of frequencies for all 3^rd^ and 4^th^ MHs from the 1000 genome database and the reason for why SNVs were removed from the final panel, if applicable.

| Location | Final Panel | Med Cov | MH >2 | SNV Len | SNV1 | SNV2 | SNV3 | Afri 3+4 | EaAs 3+4 | Euro 3+4 | AdAm 3+4 | SoAs 3+4 | AfAm 3+4 | Reason for Omitting |
| --- | --- | --- | --- | --- | --- | --- | --- | --- | --- | --- | --- | --- | --- | --- |
| chr1:23885498-23885599 |  | 692 | 25 | 102 | rs11574 | rs2067053 |  | 0.011 | 0.028 | 0.242 | 0.261 | 0.081 | 0.016 | MH>2 |
| chr1:46641827-46641837 |  | 296 | 1 | 11 | rs12030928 | rs12752892 |  | 0.017 | 0.000 | 0.125 | 0.071 | 0.080 | 0.057 | Balance |
| chr1:120057158-120057246 |  | 689 | 3 | 89 | rs6203 | rs45609334 |  | 0.033 | 0.082 | 0.235 | 0.205 | 0.157 | 0.090 | Balance |
| chr1:156846120-156846233 | Yes | 1526 | 2 | 114 | rs1800880 | rs6334 |  | 0.105 | 0.139 | 0.065 | 0.117 | 0.244 | 0.098 |  |
| chr1:226573364-226573402 | Yes | 2011 | 1 | 39 | rs1805414 | rs1805408 |  | 0.143 | 0.205 | 0.159 | 0.147 | 0.183 | 0.164 |  |
| chr1:226589833-226589958 | Yes | 361 | 2 | 126 | rs1805407 | rs1805404 |  | 0.115 | 0.251 | 0.154 | 0.147 | 0.100 | 0.164 |  |
| chr2:16042003-16042051 | Yes | 392 | 1 | 49 | rs2693006 | rs67056216 |  | 0.113 | 0.177 | 0.177 | 0.159 | 0.264 | 0.123 |  |
| chr2:16073257-16073263 | Yes | 1546 | 2 | 7 | rs12986946 | rs12986949 |  | 0.052 | 0.000 | 0.101 | 0.058 | 0.115 | 0.098 |  |
| chr2:16112814-16112828 | Yes | 835 | 1 | 15 | rs16863159 | rs6716344 |  | 0.022 | 0.276 | 0.088 | 0.244 | 0.131 | 0.148 |  |
| chr2:16113594-16113723 | Yes | 368 | 4 | 130 | rs34339850 | rs6741005 |  | 0.052 | 0.284 | 0.217 | 0.183 | 0.245 | 0.066 |  |
| chr2:29416366-29416481 |  | 677 | 2 | 116 | rs1881421 | rs1881420 |  | 0.240 | 0.000 | 0.150 | 0.127 | 0.027 | 0.312 | Balance |
| chr2:29416481-29416615 |  | 750 | 15 | 135 | rs1881420 | rs56132472 |  | 0.078 | 0.000 | 0.123 | 0.065 | 0.024 | 0.090 | MH>2 |
| chr2:29446184-29446202 | Yes | 2130 | 0 | 19 | rs2276550 | rs4622670 |  | 0.259 | 0.054 | 0.236 | 0.222 | 0.203 | 0.033 |  |
| chr2:29446701-29446721 | Yes | 686 | 1 | 21 | rs12619049 | rs4665447 |  | 0.412 | 0.081 | 0.026 | 0.062 | 0.015 | 0.320 |  |
| chr2:29447108-29447253 | Yes | 448 | 1 | 146 | rs4387740 | rs6723311 |  | 0.390 | 0.141 | 0.254 | 0.232 | 0.173 | 0.426 |  |
| chr2:47800577-47800603 | Yes | 1072 | 0 | 27 | rs56239373 | rs3814360 |  | 0.077 | 0.154 | 0.042 | 0.065 | 0.086 | 0.057 |  |
| chr2:47852559-47852643 | Yes | 293 | 5 | 85 | rs6722699 | rs10165802 |  | 0.110 | 0.076 | 0.093 | 0.104 | 0.061 | 0.139 |  |
| chr2:47856243-47856328 |  | 572 | 8 | 86 | rs11892597 | rs11887626 |  | 0.209 | 0.078 | 0.273 | 0.203 | 0.261 | 0.205 | MH>2 |
| chr2:48010488-48010558 |  | 1461 | 2 | 71 | rs1042821 | rs1042820 |  | 0.020 | 0.000 | 0.175 | 0.114 | 0.065 | 0.033 | Balance |
| chr2:112754828-112754880 |  | 366 | 1 | 53 | rs3811632 | rs3811633 |  | 0.103 | 0.106 | 0.287 | 0.189 | 0.230 | 0.156 | Balance |
| chr2:112754943-112755001 |  | 747 | 3 | 59 | rs3811634 | rs2230515 |  | 0.104 | 0.106 | 0.287 | 0.189 | 0.230 | 0.156 | Balance |
| chr2:113980645-113980736 |  | 550 | 0 | 92 | rs12472361 | rs56865224 |  | 0.017 | 0.073 | 0.110 | 0.117 | 0.129 | 0.057 |  |
| chr2:113982584-113982608 |  | 540 | 0 | 25 | rs4849177 | rs4849178 |  | 0.017 | 0.077 | 0.164 | 0.138 | 0.183 | 0.057 |  |
| chr2:113983937-113984033 | Yes | 776 | 1 | 97 | rs3748915 | rs3748916 |  | 0.203 | 0.086 | 0.163 | 0.135 | 0.229 | 0.213 |  |
| chr2:113984503-113984594 | Yes | 1400 | 0 | 92 | rs2241975 | rs67776659 |  | 0.142 | 0.013 | 0.110 | 0.087 | 0.038 | 0.074 |  |
| chr2:113985170-113985186 |  | 1536 | 2 | 17 | rs4849179 | rs4849180 |  | 0.020 | 0.077 | 0.163 | 0.138 | 0.183 | 0.066 | LD,<br>homopolymer |
| chr2:113989236-113989267 | Yes | 1009 | 2 | 32 | rs2863242 | rs2863243 |  | 0.017 | 0.074 | 0.163 | 0.138 | 0.183 | 0.057 |  |
| chr2:113990242-113990261 |  | 2923 | 0 | 20 | rs2166421 | rs7421852 |  | 0.011 | 0.074 | 0.162 | 0.135 | 0.183 | 0.041 | LD,<br>homopolymer |
| chr2:202122956-202122995 | Yes | 1337 | 0 | 40 | rs3769824 | rs3769823 |  | 0.000 | 0.000 | 0.047 | 0.114 | 0.043 | 0.016 |  |
| chr3:12641425-12641518 |  | 346 | 1 | 94 | rs5746223 | rs2246390 |  | 0.040 | 0.030 | 0.111 | 0.125 | 0.065 | 0.090 | Balance |
| chr3:12642945-12643013 |  | 277 | 3 | 69 | rs3773345 | rs904464 | rs904452 | 0.145 | 0.031 | 0.192 | 0.114 | 0.052 | 0.148 | Balance |
| chr3:12649857-12649937 | Yes | 567 | 2 | 81 | rs2055311 | rs963959 |  | 0.225 | 0.028 | 0.164 | 0.310 | 0.125 | 0.221 |  |
| chr3:36986932-36986992 | Yes | 2760 | 4 | 61 | rs2276809 | rs2276808 |  | 0.073 | 0.077 | 0.115 | 0.160 | 0.216 | 0.090 |  |
| chr3:36986992-36987083 |  | 2802 | 6 | 92 | rs2276808 | rs2276807 |  | 0.210 | 0.000 | 0.048 | 0.078 | 0.000 | 0.303 | MH>2 |
| chr3:37022803-37022864 |  | 345 | 0 | 62 | rs4678921 | rs4678922 |  | 0.082 | 0.000 | 0.000 | 0.026 | 0.000 | 0.213 | Balance |
| chr3:71247257-71247304 | Yes | 1098 | 0 | 48 | rs939845 | rs2037474 |  | 0.163 | 0.104 | 0.064 | 0.202 | 0.044 | 0.139 |  |
| chr3:138327951-138328016 | Yes | 634 | 1 | 66 | rs61699523 | rs111398337 |  | 0.167 | 0.020 | 0.028 | 0.071 | 0.110 | 0.295 |  |
| chr3:142277536-142277575 | Yes | 642 | 0 | 40 | rs2227929 | rs2227930 |  | 0.147 | 0.118 | 0.200 | 0.154 | 0.158 | 0.188 |  |
| chr3:178968634-178968660 |  | 1223 | 0 | 27 | rs7645550 | rs1170672 |  | 0.041 | 0.000 | 0.095 | 0.038 | 0.170 | 0.057 | Balance |
| chr3:178984575-178984679 | Yes | 2320 | 2 | 105 | rs7612684 | rs7646600 |  | 0.302 | 0.011 | 0.177 | 0.131 | 0.132 | 0.213 |  |
| chr3:178986121-178986203 | Yes | 623 | 5 | 83 | rs73188921 | rs9830427 | rs9830432 | 0.158 | 0.119 | 0.054 | 0.076 | 0.190 | 0.131 |  |
| chr3:178990402-178990462 | Yes | 1179 | 1 | 61 | rs2864411 | rs6443633 |  | 0.017 | 0.142 | 0.000 | 0.050 | 0.045 | 0.098 |  |
| chr3:183211906-183212026 |  | 536 | 2 | 121 | rs1520101 | rs2256061 |  | 0.128 | 0.000 | 0.182 | 0.123 | 0.162 | 0.238 | Balance |
| chr4:1745492-1745500 | Yes | 4202 | 2 | 9 | rs4865466 | rs4865467 |  | 0.126 | 0.144 | 0.217 | 0.306 | 0.229 | 0.139 |  |
| chr4:1747971-1748069 |  | 1482 | 1 | 99 | rs4572936 | rs4558926 |  | 0.049 | 0.000 | 0.156 | 0.095 | 0.082 | 0.098 | Balance |
| chr4:1748200-1748266 |  | 2245 | 8 | 67 | rs4507432 | rs4597899 | rs62285133 | 0.328 | 0.159 | 0.392 | 0.311 | 0.273 | 0.336 | MH>2 |
| chr4:1750487-1750584 | Yes | 1702 | 3 | 98 | rs7680647 | rs73202803 |  | 0.042 | 0.161 | 0.235 | 0.180 | 0.121 | 0.057 |  |
| chr4:1785738-1785791 |  | 2874 | 2 | 54 | rs1867926 | rs61407096 |  | 0.063 | 0.000 | 0.158 | 0.157 | 0.087 | 0.074 | Balance |
| chr4:1788994-1789044 | Yes | 678 | 4 | 51 | rs11248077 | rs11248078 |  | 0.249 | 0.233 | 0.383 | 0.346 | 0.377 | 0.254 |  |
| chr4:1796629-1796636 | Yes | 319 | 1 | 8 | rs3135841 | rs3135842 |  | 0.254 | 0.051 | 0.094 | 0.141 | 0.061 | 0.262 |  |
| chr4:1797741-1797852 | Yes | 995 | 4 | 112 | rs3135848 | rs743682 |  | 0.227 | 0.056 | 0.092 | 0.144 | 0.062 | 0.271 |  |
| chr4:54269096-54269173 | Yes | 557 | 1 | 78 | rs10001201 | rs62325166 |  | 0.050 | 0.133 | 0.140 | 0.105 | 0.046 | 0.082 |  |
| chr4:54526892-54526902 |  | 647 | 2 | 11 | rs12649494 | rs2668565 |  | 0.023 | 0.093 | 0.153 | 0.079 | 0.117 | 0.082 | Balance |
| chr4:54589337-54589371 |  | 445 | 21 | 35 | rs2590827 | rs75219949 |  | 0.054 | 0.223 | 0.140 | 0.208 | 0.207 | 0.115 | MH>2 |
| chr4:54657737-54657790 | Yes | 288 | 5 | 54 | rs28489910 | rs4864823 |  | 0.233 | 0.111 | 0.209 | 0.226 | 0.148 | 0.254 |  |
| chr4:55049698-55049751 |  | 310 | 3 | 54 | rs13131212 | rs7659596 |  | 0.028 | 0.061 | 0.215 | 0.159 | 0.242 | 0.041 | Balance |
| chr4:55208737-55208788 | Yes | 284 | 3 | 52 | rs2412560 | rs10018115 | rs73234206 | 0.202 | 0.247 | 0.200 | 0.270 | 0.317 | 0.361 |  |
| chr4:55477000-55477073 |  | 991 | 4 | 74 | rs17827038 | rs2703460 |  | 0.020 | 0.072 | 0.134 | 0.098 | 0.195 | 0.090 | Balance |
| chr4:55501109-55501195 | Yes | 357 | 5 | 87 | rs6554196 | rs6554197 |  | 0.110 | 0.110 | 0.200 | 0.163 | 0.223 | 0.230 |  |
| chr4:55582037-55582068 | Yes | 714 | 3 | 32 | rs76272262 | rs3134889 |  | 0.040 | 0.172 | 0.036 | 0.051 | 0.081 | 0.025 |  |
| chr4:55619846-55619859 | Yes | 892 | 3 | 14 | rs11732442 | rs4353958 |  | 0.125 | 0.109 | 0.109 | 0.069 | 0.212 | 0.139 |  |
| chr4:55916001-55916063 |  | 317 | 0 | 63 | rs7664996 | rs7665169 |  | 0.164 | 0.000 | 0.201 | 0.177 | 0.149 | 0.172 | Balance |
| chr4:55977759-55977800 |  | 278 | 3 | 42 | rs3943404 | rs2034965 |  | 0.210 | 0.215 | 0.258 | 0.264 | 0.227 | 0.213 | Balance |
| chr4:55982752-55982784 | Yes | 651 | 1 | 33 | rs11133360 | rs34945396 |  | 0.044 | 0.204 | 0.194 | 0.144 | 0.190 | 0.041 |  |
| chr4:56026865-56026914 | Yes | 565 | 1 | 50 | rs4864958 | rs75371420 | rs34743464 | 0.216 | 0.200 | 0.284 | 0.180 | 0.453 | 0.238 |  |
| chr4:106196829-106196951 | Yes | 534 | 0 | 123 | rs34402524 | rs2454206 |  | 0.066 | 0.047 | 0.140 | 0.089 | 0.090 | 0.090 |  |
| chr4:143043340-143043404 |  | 351 | 0 | 65 | rs2270658 | rs13133767 |  | 0.016 | 0.075 | 0.082 | 0.105 | 0.089 | 0.057 | Balance |
| chr4:187534362-187534375 | Yes | 2353 | 0 | 14 | rs2249916 | rs2249917 |  | 0.195 | 0.281 | 0.110 | 0.189 | 0.084 | 0.197 |  |
| chr4:187629497-187629538 | Yes | 1727 | 0 | 42 | rs458021 | rs3733413 |  | 0.128 | 0.085 | 0.070 | 0.091 | 0.031 | 0.098 |  |
| chr5:231111-231143 | Yes | 2366 | 1 | 33 | rs1126417 | rs2288459 |  | 0.164 | 0.058 | 0.111 | 0.241 | 0.079 | 0.139 |  |
| chr5:256472-256509 |  | 1282 | 9 | 38 | rs6961 | rs6962 |  | 0.269 | 0.059 | 0.105 | 0.108 | 0.075 | 0.197 | MH>2 |
| chr5:35861068-35861159 | Yes | 351 | 3 | 92 | rs1494558 | rs11567705 | rs969128 | 0.328 | 0.191 | 0.413 | 0.349 | 0.239 | 0.443 |  |
| chr5:35871190-35871273 | Yes | 255 | 1 | 84 | rs1494555 | rs2228141 |  | 0.069 | 0.153 | 0.144 | 0.166 | 0.062 | 0.164 |  |
| chr5:56178111-56178217 | Yes | 473 | 0 | 107 | rs3822625 | rs832583 |  | 0.119 | 0.108 | 0.075 | 0.078 | 0.055 | 0.049 |  |
| chr5:57754808-57754851 |  | 359 | 2 | 44 | rs697133 | rs702722 |  | 0.230 | 0.105 | 0.104 | 0.069 | 0.098 | 0.205 | Balance |
| chr5:67477132-67477234 | Yes | 371 | 0 | 103 | rs34721946 | rs34166422 | rs73126524 | 0.017 | 0.247 | 0.035 | 0.105 | 0.072 | 0.033 |  |
| chr5:67492589-67492652 | Yes | 677 | 2 | 64 | rs13188623 | rs58409263 |  | 0.105 | 0.293 | 0.121 | 0.180 | 0.118 | 0.130 |  |
| chr5:67517563-67517646 | Yes | 275 | 1 | 84 | rs6449959 | rs831227 |  | 0.243 | 0.018 | 0.187 | 0.161 | 0.100 | 0.303 |  |
| chr5:67522722-67522851 | Yes | 262 | 1 | 130 | rs706713 | rs706714 |  | 0.130 | 0.051 | 0.012 | 0.029 | 0.060 | 0.098 |  |
| chr5:67534039-67534057 | Yes | 887 | 0 | 19 | rs7709243 | rs10940158 | rs12652661 | 0.216 | 0.154 | 0.212 | 0.272 | 0.097 | 0.156 |  |
| chr5:67553771-67553827 | Yes | 584 | 1 | 57 | rs6893676 | rs34303 |  | 0.090 | 0.168 | 0.173 | 0.143 | 0.106 | 0.074 |  |
| chr5:79950497-79950512 |  | 1392 | 8 | 16 | rs2250063 | rs1105524 |  | 0.036 | 0.047 | 0.248 | 0.206 | 0.231 | 0.098 | MH>2 |
| chr5:149456772-149456811 | Yes | 1109 | 3 | 40 | rs60844779 | rs3829987 |  | 0.223 | 0.068 | 0.031 | 0.215 | 0.051 | 0.221 |  |
| chr5:176517326-176517461 |  | 652 | 3 | 136 | rs422421 | rs446382 |  | 0.169 | 0.000 | 0.078 | 0.040 | 0.033 | 0.148 | Balance |
| chr5:176523562-176523597 | Yes | 1990 | 0 | 36 | rs31777 | rs31776 |  | 0.137 | 0.000 | 0.076 | 0.038 | 0.033 | 0.115 |  |
| chr5:176531772-176531857 | Yes | 284 | 3 | 86 | rs7708357 | rs165943 |  | 0.168 | 0.046 | 0.242 | 0.248 | 0.183 | 0.197 |  |
| chr5:176539212-176539286 |  | 922 | 0 | 75 | rs244730 | rs351864 |  | 0.031 | 0.027 | 0.121 | 0.140 | 0.207 | 0.066 | Balance |
| chr5:176721198-176721272 |  | 1806 | 1 | 75 | rs28580074 | rs11740250 |  | 0.011 | 0.000 | 0.119 | 0.146 | 0.181 | 0.041 | Balance |
| chr5:180046209-180046344 |  | 765 | 12 | 136 | rs446003 | rs448012 |  | 0.100 | 0.057 | 0.083 | 0.075 | 0.135 | 0.098 | MH>2 |
| chr5:180051003-180051118 |  | 2483 | 2 | 116 | rs307826 | rs728986 |  | 0.015 | 0.000 | 0.037 | 0.055 | 0.044 | 0.016 | Balance |
| chr5:180057231-180057293 |  | 1518 | 0 | 63 | rs3736061 | rs34221241 |  | 0.000 | 0.000 | 0.081 | 0.033 | 0.041 | 0.016 | Balance |
| chr6:26056549-26056708 | Yes | 524 | 2 | 160 | rs10425 | rs2230653 | rs12204800 | 0.048 | 0.309 | 0.227 | 0.344 | 0.256 | 0.082 |  |
| chr6:30865115-30865204 | Yes | 461 | 5 | 90 | rs2239517 | rs2267641 |  | 0.120 | 0.244 | 0.038 | 0.063 | 0.094 | 0.098 |  |
| chr6:32188603-32188642 | Yes | 1185 | 1 | 40 | rs520803 | rs520692 | rs520688 | 0.000 | 0.047 | 0.000 | 0.000 | 0.011 | 0.000 |  |
| chr6:32190390-32190484 | Yes | 2363 | 5 | 95 | rs915894 | rs8192569 |  | 0.330 | 0.232 | 0.102 | 0.141 | 0.205 | 0.328 |  |
| chr6:41924853-41924931 | Yes | 922 | 2 | 79 | rs4623235 | rs16895130 |  | 0.095 | 0.110 | 0.210 | 0.156 | 0.138 | 0.107 |  |
| chr6:41941980-41941993 |  | 1079 | 1 | 14 | rs4415146 | rs72853828 |  | 0.000 | 0.089 | 0.221 | 0.125 | 0.132 | 0.049 | Balance |
| chr6:42013020-42013049 | Yes | 530 | 0 | 30 | rs9381126 | rs6919122 | rs6942118 | 0.351 | 0.421 | 0.381 | 0.504 | 0.390 | 0.254 |  |
| chr6:42039487-42039542 | Yes | 651 | 3 | 56 | rs9349215 | rs66472208 |  | 0.023 | 0.245 | 0.020 | 0.048 | 0.127 | 0.057 |  |
| chr6:42039551-42039666 | Yes | 292 | 1 | 116 | rs66489927 | rs7763360 | rs2492927 | 0.192 | 0.148 | 0.300 | 0.248 | 0.322 | 0.221 |  |
| chr6:42044204-42044273 |  | 717 | 1 | 70 | rs11968793 | rs9471750 |  | 0.184 | 0.057 | 0.046 | 0.045 | 0.014 | 0.221 | Balance |
| chr6:42052577-42052667 | Yes | 305 | 0 | 91 | rs9357387 | rs2493841 | rs9381136 | 0.050 | 0.163 | 0.176 | 0.161 | 0.139 | 0.123 |  |
| chr6:117725448-117725578 |  | 277 | 4 | 131 | rs1998206 | rs2243378 |  | 0.076 | 0.181 | 0.150 | 0.143 | 0.197 | 0.197 | Balance |
| chr6:152382311-152382325 |  | 279 | 2 | 15 | rs2273206 | rs2273207 |  | 0.137 | 0.039 | 0.026 | 0.039 | 0.055 | 0.066 | Balance |
| chr7:6026775-6026942 |  | 720 | 19 | 168 | rs2228006 | rs1805323 |  | 0.000 | 0.122 | 0.046 | 0.017 | 0.106 | 0.016 | MH>2 |
| chr7:6026942-6026988 | Yes | 3560 | 3 | 47 | rs1805323 | rs1805321 |  | 0.000 | 0.303 | 0.046 | 0.017 | 0.153 | 0.016 |  |
| chr7:55220177-55220202 | Yes | 1118 | 0 | 26 | rs11506105 | rs845561 |  | 0.115 | 0.265 | 0.254 | 0.304 | 0.413 | 0.148 |  |
| chr7:55236774-55236862 |  | 458 | 5 | 89 | rs845550 | rs1815156 |  | 0.096 | 0.000 | 0.114 | 0.062 | 0.042 | 0.123 | Balance |
| chr7:55251541-55251648 | Yes | 672 | 4 | 108 | rs2877261 | rs13222385 | rs11771471 | 0.200 | 0.076 | 0.233 | 0.183 | 0.090 | 0.189 |  |
| chr7:55306599-55306625 |  | 577 | 6 | 47 | rs2037700 | rs2037699 | rs2037698 | 0.368 | 0.083 | 0.104 | 0.124 | 0.045 | 0.271 | MH>2 |
| chr7:100410597-100410657 |  | 1469 | 8 | 61 | rs2230585 | rs770657085 |  | 0.164 | 0.056 | 0.000 | 0.043 | 0.156 | 0.197 | MH>2 |
| chr7:100416139-100416250 | Yes | 1438 | 3 | 112 | rs3857809 | rs144173 |  | 0.185 | 0.059 | 0.000 | 0.301 | 0.173 | 0.172 |  |
| chr7:116336880-116336947 | Yes | 666 | 1 | 68 | rs2237708 | rs39749 |  | 0.036 | 0.209 | 0.257 | 0.228 | 0.242 | 0.115 |  |
| chr7:116471122-116471227 | Yes | 297 | 4 | 106 | rs41773 | rs62470772 |  | 0.129 | 0.093 | 0.206 | 0.115 | 0.148 | 0.131 |  |
| chr7:116480433-116480524 |  | 319 | 7 | 92 | rs41777 | rs35099490 | rs41778 | 0.267 | 0.217 | 0.279 | 0.268 | 0.283 | 0.288 | MH>2 |
| chr8:30999122-30999123 | Yes | 554 | 3 | 2 | rs3024239 | rs2737335 |  | 0.149 | 0.024 | 0.060 | 0.032 | 0.085 | 0.246 |  |
| chr8:31024638-31024654 | Yes | 432 | 0 | 17 | rs1801196 | rs1346044 |  | 0.147 | 0.104 | 0.266 | 0.173 | 0.283 | 0.221 |  |
| chr8:38299624-38299715 | Yes | 1668 | 5 | 92 | rs60527016 | rs6987534 |  | 0.028 | 0.286 | 0.236 | 0.219 | 0.076 | 0.074 |  |
| chr8:38310910-38311001 | Yes | 1289 | 0 | 92 | rs10958700 | rs4733930 |  | 0.029 | 0.323 | 0.260 | 0.249 | 0.074 | 0.074 |  |
| chr8:38317398-38317476 |  | 1504 | 4 | 79 | rs7012413 | rs7388222 | rs3758101 | 0.213 | 0.088 | 0.076 | 0.225 | 0.115 | 0.205 | Balance |
| chr8:38350292-38350315 | Yes | 580 | 2 | 24 | rs35305468 | rs7830964 |  | 0.039 | 0.249 | 0.180 | 0.118 | 0.138 | 0.107 |  |
| chr8:38359468-38359529 |  | 1045 | 3 | 62 | rs7845393 | rs35804490 |  | 0.137 | 0.172 | 0.125 | 0.114 | 0.282 | 0.180 | LD |
| chr8:38361379-38361430 | Yes | 1456 | 2 | 52 | rs328294 | rs328293 |  | 0.309 | 0.172 | 0.126 | 0.115 | 0.283 | 0.213 |  |
| chr8:128700175-128700233 | Yes | 496 | 2 | 59 | rs13282849 | rs7005394 |  | 0.208 | 0.179 | 0.063 | 0.084 | 0.201 | 0.287 |  |
| chr8:128713221-128713364 | Yes | 796 | 5 | 144 | rs28548827 | rs7820045 |  | 0.254 | 0.057 | 0.028 | 0.101 | 0.111 | 0.148 |  |
| chr8:128718068-128718102 |  | 457 | 2 | 35 | rs9642880 | rs78635199 |  | 0.000 | 0.035 | 0.129 | 0.062 | 0.171 | 0.041 | Balance |
| chr8:128889285-128889371 | Yes | 1835 | 1 | 87 | rs6470587 | rs6470588 |  | 0.081 | 0.165 | 0.210 | 0.202 | 0.230 | 0.156 |  |
| chr8:145737636-145737816 | Yes | 485 | 0 | 181 | rs4925828 | rs4251691 |  | 0.000 | 0.203 | 0.000 | 0.072 | 0.000 | 0.000 |  |
| chr8:145741702-145741765 |  | 2058 | 6 | 64 | rs4244612 | rs4244613 |  | 0.081 | 0.027 | 0.108 | 0.094 | 0.129 | 0.148 | MH>2 |
| chr9:5408242-5408358 | Yes | 344 | 3 | 117 | rs10758685 | rs10975098 | rs10975099 | 0.084 | 0.349 | 0.257 | 0.320 | 0.409 | 0.205 |  |
| chr9:5415025-5415111 | Yes | 372 | 3 | 87 | rs78298180 | rs10758687 |  | 0.104 | 0.161 | 0.054 | 0.052 | 0.199 | 0.066 |  |
| chr9:5420254-5420266 | Yes | 1180 | 1 | 13 | rs10121219 | rs11790878 |  | 0.064 | 0.227 | 0.222 | 0.248 | 0.218 | 0.172 |  |
| chr9:5456523-5456587 |  | 277 | 1 | 65 | rs17742278 | rs7023227 | rs12002985 | 0.165 | 0.083 | 0.337 | 0.320 | 0.305 | 0.262 | Balance |
| chr9:5458035-5458095 | Yes | 323 | 3 | 61 | rs7042084 | rs10481593 |  | 0.268 | 0.132 | 0.220 | 0.249 | 0.131 | 0.205 |  |
| chr9:5484100-5484203 | Yes | 395 | 4 | 104 | rs11793113 | rs11790610 | rs10122509 | 0.139 | 0.151 | 0.094 | 0.084 | 0.167 | 0.156 |  |
| chr9:5505508-5505583 |  | 262 | 5 | 76 | rs7025653 | rs59338043 | rs6476984 | 0.228 | 0.188 | 0.234 | 0.298 | 0.215 | 0.205 | Balance |
| chr9:5514839-5514919 |  | 561 | 7 | 81 | rs2381282 | rs56381807 |  | 0.275 | 0.204 | 0.128 | 0.203 | 0.190 | 0.303 | MH>2 |
| chr9:87478135-87478172 | Yes | 1016 | 4 | 38 | rs7048015 | rs10780690 |  | 0.023 | 0.251 | 0.184 | 0.258 | 0.216 | 0.139 |  |
| chr9:93641175-93641199 |  | 693 | 2 | 25 | rs2306041 | rs2306040 |  | 0.062 | 0.000 | 0.064 | 0.058 | 0.020 | 0.066 | Balance |
| chr9:98238358-98238379 | Yes | 3840 | 0 | 22 | rs2066836 | rs1805155 |  | 0.011 | 0.083 | 0.109 | 0.076 | 0.060 | 0.074 |  |
| chr9:139401504-139401577 | Yes | 1346 | 1 | 74 | rs3124596 | rs7870145 | rs3829116 | 0.310 | 0.000 | 0.163 | 0.117 | 0.264 | 0.238 |  |
| chr9:139403268-139403280 |  | 500 | 1 | 13 | rs3125000 | rs11145765 |  | 0.046 | 0.000 | 0.095 | 0.059 | 0.238 | 0.025 | Balance |
| chr9:139405093-139405261 | Yes | 626 | 3 | 169 | rs36119806 | rs3125001 |  | 0.150 | 0.012 | 0.102 | 0.065 | 0.184 | 0.156 |  |
| chr9:139410424-139410589 | Yes | 327 | 2 | 166 | rs3125006 | rs4880099 |  | 0.088 | 0.052 | 0.115 | 0.068 | 0.215 | 0.049 |  |
| chr9:139411714-139411880 |  | 428 | 5 | 167 | rs11145767 | rs9411254 |  | 0.209 | 0.000 | 0.000 | 0.025 | 0.000 | 0.213 | Balance |
| chr10:43611708-43611865 | Yes | 629 | 2 | 158 | rs741968 | rs2256550 |  | 0.060 | 0.218 | 0.161 | 0.212 | 0.284 | 0.139 |  |
| chr10:43615505-43615633 | Yes | 463 | 5 | 129 | rs2472737 | rs1800863 |  | 0.105 | 0.121 | 0.193 | 0.187 | 0.160 | 0.115 |  |
| chr10:70332580-70332672 | Yes | 549 | 1 | 93 | rs10823229 | rs12773594 |  | 0.023 | 0.173 | 0.185 | 0.151 | 0.271 | 0.090 |  |
| chr10:104386934-104387019 | Yes | 250 | 0 | 86 | rs17114803 | rs12414407 |  | 0.224 | 0.250 | 0.093 | 0.238 | 0.240 | 0.197 |  |
| chr10:123194558-123194609 | Yes | 384 | 0 | 52 | rs7911440 | rs6585731 |  | 0.051 | 0.211 | 0.242 | 0.082 | 0.243 | 0.107 |  |
| chr10:123199092-123199095 | Yes | 1151 | 2 | 4 | rs4752560 | rs2114689 |  | 0.283 | 0.023 | 0.075 | 0.156 | 0.160 | 0.131 |  |
| chr10:123242724-123242780 |  | 415 | 2 | 57 | rs3135811 | rs3135810 |  | 0.154 | 0.000 | 0.064 | 0.030 | 0.000 | 0.148 | Balance |
| chr10:123275662-123275666 | Yes | 320 | 1 | 5 | rs2912761 | rs2981453 |  | 0.211 | 0.000 | 0.000 | 0.050 | 0.000 | 0.197 |  |
| chr10:123335839-123335866 | Yes | 1055 | 1 | 28 | rs45631611 | rs10886946 |  | 0.017 | 0.113 | 0.071 | 0.055 | 0.114 | 0.041 |  |
| chr10:123346116-123346190 | Yes | 420 | 0 | 75 | rs2981575 | rs1219648 |  | 0.195 | 0.048 | 0.000 | 0.022 | 0.013 | 0.090 |  |
| chr10:123396636-123396715 |  | 259 | 5 | 80 | rs10788193 | rs2454809 |  | 0.059 | 0.176 | 0.195 | 0.192 | 0.246 | 0.139 | Coverage |
| chr10:123396728-123396806 | Yes | 331 | 2 | 79 | rs1909670 | rs1614303 |  | 0.029 | 0.176 | 0.100 | 0.131 | 0.073 | 0.057 |  |
| chr10:123406645-123406663 | Yes | 699 | 4 | 19 | rs10788194 | rs7923788 |  | 0.084 | 0.227 | 0.151 | 0.192 | 0.125 | 0.139 |  |
| chr11:534197-534242 | Yes | 2026 | 1 | 46 | rs41258054 | rs12628 |  | 0.000 | 0.153 | 0.056 | 0.137 | 0.076 | 0.025 |  |
| chr11:8246326-8246343 |  | 287 | 6 | 18 | rs34544683 | rs3816490 |  | 0.022 | 0.098 | 0.125 | 0.219 | 0.133 | 0.123 | MH>2 |
| chr11:69412090-69412124 | Yes | 2968 | 1 | 35 | rs79274134 | rs7112989 |  | 0.254 | 0.232 | 0.000 | 0.127 | 0.031 | 0.254 |  |
| chr11:69446719-69446766 |  | 1480 | 2 | 48 | rs609581 | rs11826558 |  | 0.070 | 0.015 | 0.058 | 0.079 | 0.187 | 0.139 | Balance |
| chr11:69448373-69448445 |  | 563 | 7 | 73 | rs654240 | rs654648 |  | 0.188 | 0.029 | 0.136 | 0.112 | 0.210 | 0.221 | MH>2 |
| chr11:69508082-69508159 |  | 574 | 3 | 78 | rs7930020 | rs10908198 |  | 0.000 | 0.041 | 0.125 | 0.267 | 0.045 | 0.066 | Balance |
| chr12:4333159-4333203 |  | 1194 | 4 | 45 | rs7955734 | rs7970219 | rs7954847 | 0.268 | 0.092 | 0.026 | 0.085 | 0.100 | 0.229 | Balance |
| chr12:4346169-4346177 | Yes | 646 | 0 | 9 | rs11063052 | rs11832328 |  | 0.318 | 0.079 | 0.038 | 0.072 | 0.080 | 0.213 |  |
| chr12:4351884-4352027 | Yes | 468 | 5 | 144 | rs7955545 | rs4766223 |  | 0.051 | 0.113 | 0.033 | 0.076 | 0.092 | 0.139 |  |
| chr12:4376089-4376091 | Yes | 306 | 2 | 3 | rs4238013 | rs12818766 |  | 0.119 | 0.033 | 0.181 | 0.161 | 0.147 | 0.139 |  |
| chr12:4399036-4399054 | Yes | 1619 | 2 | 52 | rs3217859 | rs3217860 | rs3217861 | 0.325 | 0.391 | 0.414 | 0.491 | 0.479 | 0.303 |  |
| chr12:4399917-4399970 | Yes | 892 | 2 | 54 | rs3217867 | rs3217868 | rs3217869 | 0.173 | 0.041 | 0.220 | 0.133 | 0.188 | 0.213 |  |
| chr12:4411639-4411683 | Yes | 1376 | 1 | 45 | rs3217925 | rs3217926 |  | 0.127 | 0.068 | 0.253 | 0.172 | 0.227 | 0.188 |  |
| chr12:4417127-4417232 | Yes | 1224 | 1 | 106 | rs7133323 | rs9668504 |  | 0.449 | 0.324 | 0.237 | 0.282 | 0.142 | 0.418 |  |
| chr12:4463020-4463036 |  | 850 | 31 | 17 | rs7297048 | rs7296130 | rs12823973 | 0.115 | 0.089 | 0.193 | 0.303 | 0.117 | 0.131 | MH>2 |
| chr12:12009741-12009874 | Yes | 379 | 2 | 134 | rs2238126 | rs743614 |  | 0.181 | 0.240 | 0.190 | 0.249 | 0.079 | 0.180 |  |
| chr12:12013572-12013612 | Yes | 647 | 3 | 41 | rs2855708 | rs6488463 |  | 0.232 | 0.196 | 0.211 | 0.347 | 0.146 | 0.262 |  |
| chr12:12016008-12016089 | Yes | 1488 | 3 | 82 | rs2238130 | rs2416944 | rs2238131 | 0.125 | 0.248 | 0.144 | 0.216 | 0.104 | 0.197 |  |
| chr12:12020114-12020170 | Yes | 637 | 1 | 57 | rs2723805 | rs7973930 |  | 0.241 | 0.111 | 0.075 | 0.066 | 0.054 | 0.238 |  |
| chr12:12033536-12033570 |  | 2184 | 4 | 35 | rs11832110 | rs4763730 |  | 0.232 | 0.000 | 0.042 | 0.074 | 0.000 | 0.189 | Balance |
| chr12:12035649-12035664 | Yes | 2052 | 1 | 16 | rs2710310 | rs2739085 |  | 0.126 | 0.271 | 0.194 | 0.251 | 0.159 | 0.180 |  |
| chr12:18656174-18656225 |  | 381 | 1 | 52 | rs11044141 | rs11044142 |  | 0.099 | 0.000 | 0.000 | 0.000 | 0.000 | 0.041 | Balance |
| chr12:56494991-56494998 |  | 3387 | 6 | 8 | rs2271189 | rs773123 |  | 0.073 | 0.000 | 0.110 | 0.066 | 0.070 | 0.066 | MH>2 |
| chr12:69169222-69169316 | Yes | 404 | 3 | 95 | rs6581833 | rs73334654 |  | 0.256 | 0.016 | 0.059 | 0.078 | 0.000 | 0.328 |  |
| chr12:69265196-69265278 | Yes | 768 | 0 | 83 | rs3817605 | rs2293637 |  | 0.310 | 0.192 | 0.022 | 0.111 | 0.106 | 0.271 |  |
| chr12:69277127-69277165 | Yes | 773 | 1 | 39 | rs10878875 | rs1663588 |  | 0.126 | 0.162 | 0.124 | 0.133 | 0.215 | 0.180 |  |
| chr12:121416622-121416650 | Yes | 3076 | 2 | 29 | rs1169289 | rs1169288 |  | 0.082 | 0.049 | 0.132 | 0.112 | 0.151 | 0.131 |  |
| chr12:121431272-121431300 | Yes | 1774 | 0 | 29 | rs2071190 | rs1169301 |  | 0.118 | 0.255 | 0.236 | 0.272 | 0.182 | 0.189 |  |
| chr12:121435427-121435475 |  | 3503 | 1 | 49 | rs2464196 | rs2464195 |  | 0.014 | 0.000 | 0.062 | 0.046 | 0.092 | 0.049 | Balance |
| chr12:121437114-121437221 |  | 1919 | 0 | 108 | rs55834942 | rs1169304 |  | 0.012 | 0.000 | 0.166 | 0.082 | 0.023 | 0.025 | Balance |
| chr12:133208886-133208979 | Yes | 739 | 2 | 94 | rs5745023 | rs5745022 |  | 0.173 | 0.105 | 0.135 | 0.219 | 0.049 | 0.180 |  |
| chr12:133226159-133226196 | Yes | 587 | 2 | 38 | rs4883613 | rs4883537 |  | 0.105 | 0.107 | 0.135 | 0.222 | 0.050 | 0.180 |  |
| chr12:133253995-133254083 | Yes | 448 | 1 | 89 | rs5744751 | rs5744750 |  | 0.000 | 0.105 | 0.100 | 0.045 | 0.042 | 0.025 |  |
| chr13:21562832-21562948 |  | 1715 | 3 | 117 | rs2770928 | rs558614 |  | 0.175 | 0.000 | 0.080 | 0.087 | 0.153 | 0.156 | Balance |
| chr13:32854998-32855018 |  | 333 | 1 | 21 | rs17692070 | rs11617975 |  | 0.129 | 0.000 | 0.098 | 0.128 | 0.122 | 0.123 | Balance |
| chr13:32986219-32986340 | Yes | 313 | 0 | 122 | rs206319 | rs206320 | rs615762 | 0.107 | 0.204 | 0.175 | 0.244 | 0.262 | 0.107 |  |
| chr14:35872792-35872926 |  | 643 | 1 | 135 | rs2233415 | rs1050851 |  | 0.020 | 0.019 | 0.213 | 0.130 | 0.146 | 0.033 | Balance |
| chr14:102568296-102568367 |  | 969 | 0 | 72 | rs10873531 | rs8005905 |  | 0.278 | 0.049 | 0.017 | 0.068 | 0.123 | 0.303 | Balance |
| chr14:104165753-104165927 |  | 765 | 4 | 175 | rs861539 | rs1799796 |  | 0.114 | 0.073 | 0.295 | 0.229 | 0.189 | 0.115 | Balance |
| chr14:105239146-105239192 | Yes | 521 | 5 | 47 | rs3803304 | rs2494732 |  | 0.169 | 0.097 | 0.171 | 0.290 | 0.302 | 0.230 |  |
| chr14:105258892-105258893 | Yes | 737 | 1 | 2 | rs2494748 | rs2494749 |  | 0.120 | 0.122 | 0.092 | 0.231 | 0.245 | 0.123 |  |
| chr15:41857216-41857303 |  | 1528 | 2 | 88 | rs11639399 | rs2277536 |  | 0.096 | 0.012 | 0.308 | 0.172 | 0.236 | 0.139 | Balance |
| chr15:41860411-41860490 |  | 860 | 2 | 80 | rs7171675 | rs12148316 |  | 0.095 | 0.011 | 0.134 | 0.131 | 0.110 | 0.139 | Balance |
| chr15:67457335-67457485 | Yes | 475 | 4 | 151 | rs1065080 | rs2289261 |  | 0.133 | 0.238 | 0.139 | 0.087 | 0.220 | 0.156 |  |
| chr15:73996066-73996101 |  | 3299 | 3 | 36 | rs11574483 | rs10083681 |  | 0.147 | 0.000 | 0.000 | 0.017 | 0.000 | 0.148 | Balance |
| chr15:88488326-88488428 | Yes | 1800 | 1 | 103 | rs8042993 | rs1369426 |  | 0.088 | 0.135 | 0.153 | 0.097 | 0.261 | 0.066 |  |
| chr15:88549118-88549151 | Yes | 1763 | 0 | 34 | rs11073758 | rs12324332 |  | 0.266 | 0.015 | 0.124 | 0.133 | 0.079 | 0.303 |  |
| chr15:88554878-88554886 |  | 2501 | 3 | 9 | rs4989257 | rs898707 |  | 0.040 | 0.014 | 0.124 | 0.128 | 0.082 | 0.164 | Balance |
| chr15:88646922-88647038 | Yes | 975 | 1 | 117 | rs16941255 | rs76506232 |  | 0.110 | 0.132 | 0.000 | 0.010 | 0.000 | 0.115 |  |
| chr15:88667852-88667948 | Yes | 1099 | 0 | 97 | rs3784411 | rs3784410 |  | 0.192 | 0.100 | 0.217 | 0.225 | 0.151 | 0.172 |  |
| chr16:2138218-2138269 |  | 1617 | 14 | 52 | rs1800718 | rs1748 |  | 0.136 | 0.000 | 0.090 | 0.063 | 0.024 | 0.074 | MH>2 |
| chr16:2138269-2138398 |  | 941 | 4 | 130 | rs1748 | rs13332221 |  | 0.249 | 0.000 | 0.116 | 0.017 | 0.123 | 0.197 | Balance |
| chr16:2138398-2138422 | Yes | 2026 | 0 | 25 | rs13332221 | rs13332222 |  | 0.118 | 0.000 | 0.000 | 0.013 | 0.000 | 0.090 |  |
| chr16:81819768-81819820 | Yes | 2558 | 1 | 53 | rs1143685 | rs4294811 |  | 0.140 | 0.141 | 0.282 | 0.271 | 0.126 | 0.148 |  |
| chr16:89806343-89806347 | Yes | 601 | 2 | 5 | rs11647746 | rs7195906 |  | 0.161 | 0.013 | 0.074 | 0.035 | 0.134 | 0.156 |  |
| chr16:89849480-89849629 |  | 275 | 2 | 150 | rs2239359 | rs12448860 |  | 0.032 | 0.013 | 0.064 | 0.370 | 0.352 | 0.319 | Balance |
| chr16:89858505-89858525 |  | 698 | 3 | 21 | rs6500452 | rs1800287 |  | 0.177 | 0.012 | 0.073 | 0.043 | 0.133 | 0.221 | Balance |
| chr17:1782952-1782957 | Yes | 1284 | 1 | 6 | rs5030755 | rs2230930 |  | 0.000 | 0.000 | 0.102 | 0.020 | 0.024 | 0.016 |  |
| chr17:37807698-37807707 |  | 1082 | 2 | 10 | rs9972882 | rs75397452 |  | 0.141 | 0.000 | 0.047 | 0.042 | 0.000 | 0.131 | Balance |
| chr17:37832279-37832315 | Yes | 1408 | 1 | 37 | rs1495100 | rs2934953 |  | 0.194 | 0.000 | 0.016 | 0.062 | 0.053 | 0.188 |  |
| chr17:37833562-37833567 |  | 2351 | 2 | 6 | rs11656146 | rs907088 |  | 0.000 | 0.000 | 0.092 | 0.030 | 0.011 | 0.016 | Balance |
| chr17:37834357-37834367 |  | 666 | 0 | 11 | rs732084 | rs732083 |  | 0.181 | 0.000 | 0.000 | 0.043 | 0.011 | 0.156 | Balance |
| chr17:37834715-37834808 | Yes | 1558 | 5 | 94 | rs12150603 | rs72832915 |  | 0.042 | 0.153 | 0.308 | 0.196 | 0.235 | 0.131 |  |
| chr17:37853048-37853097 |  | 1655 | 1 | 50 | rs2643194 | rs2517951 |  | 0.188 | 0.000 | 0.017 | 0.085 | 0.081 | 0.303 | Balance |
| chr17:37893458-37893484 |  | 499 | 8 | 27 | rs67597968 | rs4795393 |  | 0.209 | 0.000 | 0.053 | 0.069 | 0.098 | 0.221 | MH>2 |
| chr17:41616392-41616456 | Yes | 1646 | 1 | 65 | rs76280498 | rs7222604 |  | 0.000 | 0.150 | 0.106 | 0.110 | 0.181 | 0.033 |  |
| chr17:78820329-78820374 | Yes | 3252 | 0 | 46 | rs3751945 | rs2589156 |  | 0.082 | 0.000 | 0.107 | 0.078 | 0.115 | 0.066 |  |
| chr17:78865546-78865630 | Yes | 631 | 3 | 85 | rs2289764 | rs2289765 |  | 0.289 | 0.044 | 0.111 | 0.110 | 0.115 | 0.303 |  |
| chr17:78896488-78896529 | Yes | 2726 | 4 | 42 | rs2271602 | rs2271603 |  | 0.154 | 0.196 | 0.321 | 0.291 | 0.307 | 0.197 |  |
| chr17:78897547-78897561 | Yes | 1725 | 0 | 15 | rs7217786 | rs6565491 |  | 0.031 | 0.199 | 0.122 | 0.111 | 0.249 | 0.049 |  |
| chr17:78921117-78921211 | Yes | 1576 | 2 | 95 | rs4969231 | rs9912373 |  | 0.022 | 0.079 | 0.124 | 0.114 | 0.060 | 0.082 |  |
| chr19:2226676-2226772 | Yes | 2349 | 1 | 97 | rs3815308 | rs2302061 |  | 0.034 | 0.182 | 0.143 | 0.172 | 0.203 | 0.090 |  |
| chr19:3110349-3110361 |  | 2345 | 5 | 13 | rs11085000 | rs41276846 |  | 0.018 | 0.193 | 0.112 | 0.063 | 0.188 | 0.016 |  |
| chr19:3119184-3119239 | Yes | 1438 | 1 | 56 | rs308046 | rs4900 |  | 0.166 | 0.233 | 0.135 | 0.101 | 0.275 | 0.271 |  |
| chr19:3119405-3119406 |  | 884 | 33 | 2 | rs370286 | rs451778 |  | 0.000 | 0.233 | 0.135 | 0.087 | 0.280 | 0.025 | MH>2 |
| chr19:5210622-5210782 |  | 740 | 2 | 161 | rs2302224 | rs1143698 |  | 0.166 | 0.066 | 0.126 | 0.134 | 0.090 | 0.131 | Balance |
| chr19:5210762-5210782 |  | 4185 | 0 | 21 | rs1143699 | rs1143698 |  | 0.222 | 0.000 | 0.099 | 0.081 | 0.056 | 0.205 | Balance |
| chr19:5212380-5212482 |  | 1945 | 1 | 103 | rs1064300 | rs2230611 |  | 0.115 | 0.000 | 0.124 | 0.135 | 0.126 | 0.098 | Balance |
| chr19:7163154-7163230 | Yes | 810 | 2 | 77 | rs2963 | rs2245648 |  | 0.186 | 0.025 | 0.065 | 0.068 | 0.141 | 0.148 |  |
| chr19:7166376-7166388 | Yes | 1028 | 2 | 13 | rs2059806 | rs2229429 |  | 0.179 | 0.065 | 0.191 | 0.144 | 0.262 | 0.197 |  |
| chr19:10267011-10267077 | Yes | 265 | 0 | 67 | rs4804490 | rs2228611 |  | 0.171 | 0.281 | 0.068 | 0.184 | 0.224 | 0.172 |  |
| chr19:17937758-17937786 |  | 1721 | 0 | 29 | rs3212798 | rs3212797 |  | 0.074 | 0.000 | 0.052 | 0.033 | 0.012 | 0.066 | Balance |
| chr19:17955001-17955021 | Yes | 1946 | 1 | 21 | rs3212713 | rs3212712 | rs3212711 | 0.197 | 0.000 | 0.000 | 0.022 | 0.000 | 0.139 |  |
| chr19:30253901-30253998 | Yes | 768 | 2 | 98 | rs117342492 | rs4805475 |  | 0.000 | 0.221 | 0.000 | 0.104 | 0.073 | 0.033 |  |
| chr19:30255068-30255090 | Yes | 495 | 2 | 23 | rs8103966 | rs8099838 |  | 0.043 | 0.310 | 0.250 | 0.232 | 0.252 | 0.082 |  |
| chr19:30290349-30290357 | Yes | 2732 | 1 | 9 | rs1473201 | rs111640872 |  | 0.085 | 0.106 | 0.247 | 0.180 | 0.213 | 0.066 |  |
| chr19:30319892-30319896 |  | 284 | 5 | 5 | rs56285422 | rs8106054 |  | 0.085 | 0.106 | 0.248 | 0.180 | 0.213 | 0.066 | LD |
| chr19:30340381-30340412 | Yes | 593 | 3 | 32 | rs929813 | rs929814 |  | 0.216 | 0.087 | 0.121 | 0.293 | 0.263 | 0.271 |  |
| chr19:30361995-30362112 | Yes | 290 | 2 | 118 | rs255270 | rs255271 |  | 0.184 | 0.104 | 0.037 | 0.068 | 0.012 | 0.271 |  |
| chr19:41724820-41724885 | Yes | 2049 | 0 | 66 | rs2301236 | rs28364580 |  | 0.094 | 0.179 | 0.224 | 0.148 | 0.275 | 0.172 |  |
| chr19:41781493-41781579 | Yes | 1040 | 2 | 87 | rs8103839 | rs9304592 |  | 0.067 | 0.073 | 0.000 | 0.066 | 0.064 | 0.066 |  |
| chr19:50919797-50919828 | Yes | 2886 | 5 | 32 | rs3218776 | rs3218760 |  | 0.125 | 0.139 | 0.075 | 0.148 | 0.275 | 0.156 |  |
| chr20:9543622-9543681 | Yes | 813 | 5 | 60 | rs2297345 | rs2297346 |  | 0.122 | 0.214 | 0.088 | 0.174 | 0.059 | 0.139 |  |
| chr20:30729488-30729523 | Yes | 3150 | 2 | 36 | rs6089193 | rs6089194 |  | 0.206 | 0.085 | 0.026 | 0.137 | 0.053 | 0.189 |  |
| chr20:40714479-40714540 | Yes | 1095 | 1 | 62 | rs2016647 | rs1569548 |  | 0.114 | 0.074 | 0.242 | 0.167 | 0.138 | 0.066 |  |
| chr20:40714539-40714540 |  | 1134 | 12 | 2 | rs1569547 | rs1569548 |  | 0.000 | 0.073 | 0.231 | 0.150 | 0.120 | 0.016 | MH>2 |
| chr20:57478807-57478939 |  | 711 | 8 | 133 | rs7121 | rs3730168 |  | 0.186 | 0.091 | 0.286 | 0.120 | 0.169 | 0.205 | MH>2 |
| chr21:42845374-42845383 | Yes | 6069 | 0 | 10 | rs2298659 | rs17854725 |  | 0.173 | 0.115 | 0.230 | 0.218 | 0.189 | 0.220 |  |
| chr21:42876400-42876447 | Yes | 2128 | 0 | 48 | rs7277080 | rs395584 |  | 0.287 | 0.017 | 0.019 | 0.235 | 0.212 | 0.229 |  |
| chr21:45643008-45643052 |  | 288 | 31 | 45 | rs1048710 | rs2298564 |  | 0.069 | 0.237 | 0.045 | 0.140 | 0.043 | 0.066 | MH>2 |
| chr22:17640022-17640045 | Yes | 1258 | 0 | 24 | rs11550530 | rs7287672 |  | 0.125 | 0.035 | 0.086 | 0.130 | 0.058 | 0.066 |  |
| chr22:21337266-21337325 | Yes | 565 | 4 | 60 | rs178280 | rs13054014 |  | 0.116 | 0.200 | 0.259 | 0.223 | 0.234 | 0.189 |  |
| chr22:21348914-21349037 |  | 1246 | 25 | 124 | rs4822790 | rs178292 |  | 0.105 | 0.224 | 0.135 | 0.112 | 0.142 | 0.139 | MH>2 |
| chr22:24158895-24158899 | Yes | 713 | 2 | 5 | rs9608192 | rs2070457 |  | 0.098 | 0.059 | 0.115 | 0.071 | 0.153 | 0.090 |  |
| chr22:29690246-29690345 | Yes | 259 | 0 | 100 | rs73156524 | rs131189 |  | 0.032 | 0.281 | 0.086 | 0.053 | 0.034 | 0.090 |  |
| Final Panel | 164 | 768 | 1.78 | 59 |  |  |  | 0.139 | 0.139 | 0.139 | 0.156 | 0.153 | 0.155 |  |

**Supplementary Table 2.** Detection of in silico mixtures versus 3^rd^ MH threshold. Pure samples were mixed in 15 different combinations from each ancestry group (Afri, EaAs, Euro) as well as 15 different mixed ancestry combinations. For each threshold 3^rd^ MH frequency, the percent of all samples that scored as contaminated at that level of contamination are listed at each level of in silico mixing.

|  |  | Threshold Median 3rd MH Frequency |  |  |  |  |
| --- | --- | --- | --- | --- | --- | --- |
|  |  | 0.5 | 1 | 1.5 | 2 | 2.5 |
| In silico<br>Mixing<br>Levels | 0.5 | 40 | 0 | 0 | 0 | 0 |
|  | 1 | 100 | 0 | 0 | 0 | 0 |
|  | 1.5 | 100 | 7 | 0 | 0 | 0 |
|  | 2 | 100 | 63 | 0 | 0 | 0 |
|  | 2.5 | 100 | 100 | 2 | 0 | 0 |
|  | 3 | 100 | 100 | 35 | 0 | 0 |
|  | 4 | 100 | 100 | 95 | 15 | 0 |
|  | 5 | 100 | 100 | 100 | 88 | 7 |
|  | 10 | 100 | 100 | 100 | 100 | 100 |

